# The *Vibrio cholerae* quorum-sensing protein VqmA integrates cell density, environmental, and host-derived cues into the control of virulence

**DOI:** 10.1101/2020.05.04.076810

**Authors:** Ameya A. Mashruwala, Bonnie L. Bassler

## Abstract

Quorum sensing is a chemical communication process in which bacteria use the production, release, and detection of signal molecules called autoinducers to orchestrate collective behaviors. The human pathogen *Vibrio cholerae* requires quorum sensing to infect the small intestine. There, *V. cholerae* encounters the absence of oxygen and the presence of bile. We show that these two stimuli differentially affect quorum sensing function and, in turn, *V. cholerae* pathogenicity. The quorum-sensing receptor-transcription factor called VqmA, that detects the autoinducer called DPO, also detects the lack of oxygen and the presence of bile. Detection occurs via DPO-, oxygen-, bile-, and redox-responsive disulfide bonds that alter VqmA DNA binding activity. We propose that VqmA serves as an information processing hub that integrates quorum-sensing information, redox status, the presence or absence of oxygen, and host cues. In response to the information acquired through this mechanism, *V. cholerae* appropriately modulates its virulence output.

**Lay Abstract:** Quorum sensing (QS) is a process of chemical communication bacteria use to orchestrate collective behaviors. QS communication relies on chemical signal molecules called autoinducers. QS regulates virulence in *Vibrio cholerae*, the causative agent of the disease cholera. Transit into the human small intestine, the site of cholera infection, exposes *V. cholerae* to the host environment. In this study, we show that the combination of two stimuli encountered in the small intestine, the absence of oxygen and the presence of host-produced bile, impinge on *V. cholerae* QS function and, in turn, pathogenicity. We suggest that possessing a QS system that is responsive to multiple environmental, host, and cell density cues enables *V. cholerae* to fine-tune its virulence capacity in the human intestine.

## Background

Quorum sensing (QS) is a process of cell-cell communication that bacteria use to synchronize group behaviors such as bioluminescence, DNA exchange, virulence factor production, and biofilm formation (20, 39, 45, 56). QS depends on the production, release, accumulation, and group-wide detection of extracellular signaling molecules called autoinducers (AI) (45, 53). At low cell density (LCD), when there are few cells present and the concentration of AIs is low, the expression of genes driving individual behaviors occurs (45, 53, 59). As the cells grow to high cell density (HCD), the extracellular concentration of AIs likewise increases. Detection of accumulated AIs drives the population-wide expression of genes required for group behaviors.

*Vibrio cholerae* is a Gram-negative enteric pathogen that causes infectious gastroenteritis. In *V. cholerae*, QS regulates collective behaviors including virulence factor production and biofilm formation (27, 39, 66, 67). Specifically, at LCD, genes encoding virulence factors and those required for biofilm formation are expressed (39). At HCD, genes required for both of these traits are repressed by QS (39). This pattern of gene expression is best understood in the context of the cholera disease. Infection is initiated by the ingestion of a small number of *V. cholerae* cells, and biofilm formation and virulence factor production are required for successful colonization (66, 67). In the host, the growth-dependent accumulation of AIs launches the HCD QS program, which suppresses virulence factor production and biofilm formation, and triggers dispersal of the bacteria back into the environment. Indeed, *V. cholerae* strains “locked” into the LCD QS mode are more proficient in host colonization than strains “locked” in the HCD QS mode (27). Thus, QS is proposed to be crucial for *V. cholerae* transitions between environmental reservoirs and human hosts.

*V. cholerae* produces and detects three AIs, called AI-2, CAI-1and DPO (Figure 1) (21, 39, 47, 54). CAI-1 is used for intra-genus communication while AI-2 and DPO are employed for inter-species communication (5, 39, 47). Different combinations of the three AIs are thought to allow *V. cholerae* to distinguish the number of vibrio cells present relative to the total bacterial consortium. *V. cholerae* uses the information encoded in blends of AIs to tailor its QS output depending on whether vibrios are in the minority or the majority of a mixed-species population (5, 39).

**Figure 1.**
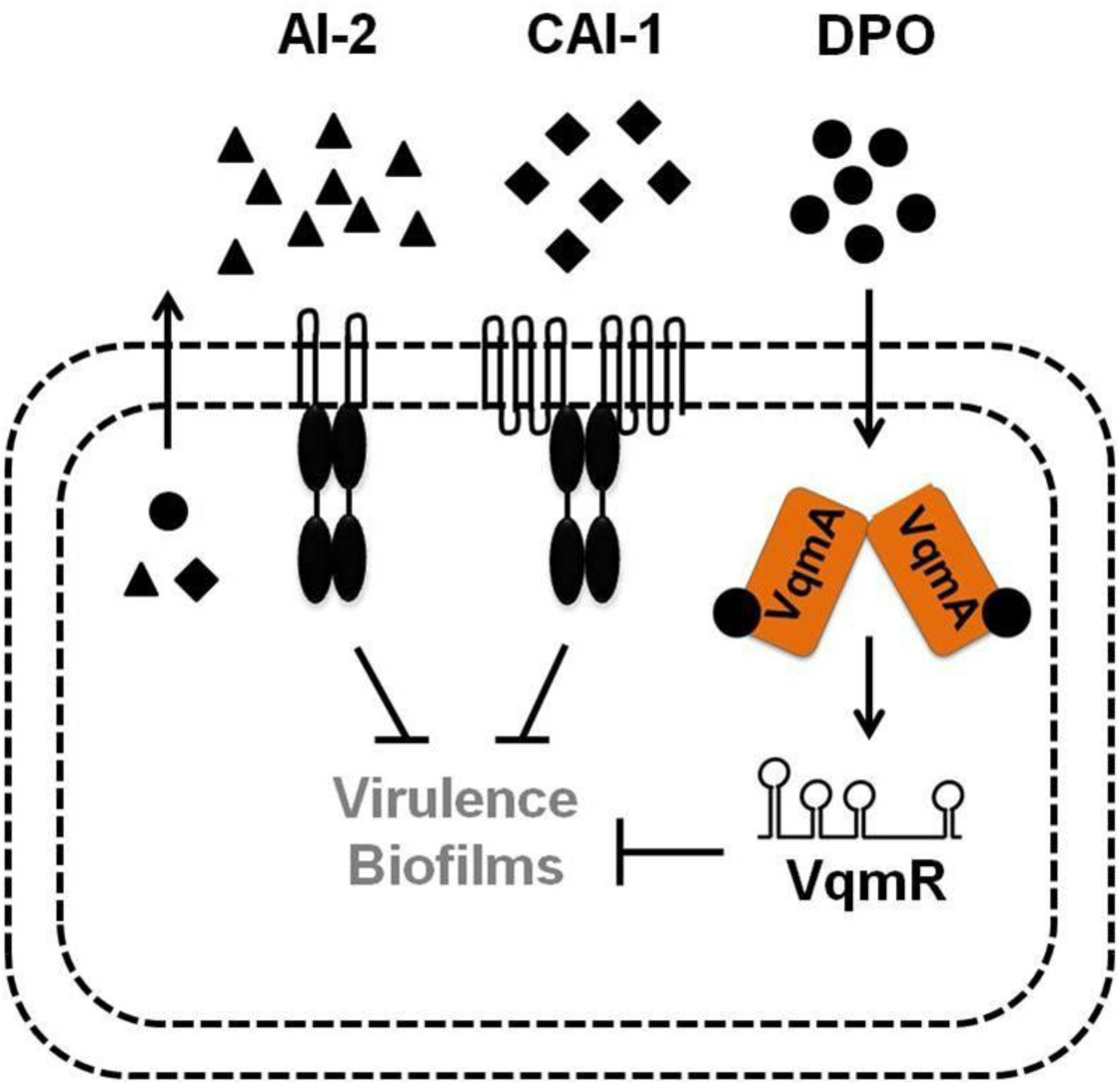
Simplified *V. cholerae* quorum-sensing pathways. See text for details.

AI-2 and CAI-1 are detected by membrane-bound receptors that funnel information into a shared regulatory pathway (Figure 1) (21, 39). DPO is detected by the cytoplasmic VqmA receptor-transcription factor that activates expression of *vqmR*, encoding the VqmR regulatory small RNA (sRNA) (34, 46, 47). Both Apo- and DPO-bound Holo-VqmA can activate *vqmR* expression with Holo-VqmA being more potent than Apo-VqmA (22). VqmR post-transcriptionally regulates target mRNAs (46). Important to this study is that at HCD, all three QS systems repress genes required for virulence and biofilm formation (Figure 1).

Upon the transition from the marine niche to the human host, *V. cholerae* switches from an aerobic to an anaerobic environment (15, 64). In addition, it encounters bile, which is abundant in the lower intestine, the primary site of *V. cholerae* infection. Bile is a heterogeneous mixture of compounds, including electrolytes and bile acids, and is estimated to be present at ∼0.2-2% weight/volume of intestinal contents (14). Bile is known to affect *V. cholerae* virulence gene expression by modulating activities of the oxidoreductase DsbA, the transmembrane-spanning transcription factor TcpP, and the ToxT transcription factor (6, 9, 23, 60, 62, 63). Bile also promotes biofilm formation in *V. cholerae*, and the second messenger molecule called cyclic-di-guanylate is involved in mediating this effect (24, 29).

Here, first we explore whether oxygen levels modulate QS in *V. cholerae*. We find that *V. cholerae* cultured under anaerobic conditions does not produce CAI-1, whereas increased DPO production does occur. In this work we focus on DPO. We show that the VqmA-DPO complex more strongly activates target gene expression under anaerobic than aerobic conditions. One consequence of the presence/absence of oxygen is an altered reducing/oxidizing (hereafter, redox) cellular environment. We show that oxygen-dependent changes in VqmA activity are governed by cysteine disulfide bonds that are responsive to the redox environment. In the absence of DPO, during aerobic growth, Apo-VqmA forms an intra-molecular disulfide-bond that limits VqmA activity. By contrast, DPO-bound VqmA forms an inter-molecular disulfide-bond that enhances VqmA activity. The formation of the inter-molecular bond is not affected by oxygen levels. In the small intestine, *V. cholerae* encounters both the absence of oxygen and the presence of bile. Bile salts inhibit formation of the inter-molecular disulfide bond in VqmA. Thus, bile and DPO have opposing effects on VqmA-DPO activity. We propose that the VqmA-DPO-VqmR QS pathway allows *V. cholerae* to integrate QS information, host cues, and environmental stimuli into the control of genes required for transitions between the human host and the environment.

## Results

### *V. cholerae* does not produce the CAI-1 QS AI under anaerobic conditions

To our knowledge, *V. cholerae* QS has only been studied under aerobic conditions. We know that the marine-human host lifecycle demands that *V. cholerae* transition between environments containing widely varying oxygen levels (15, 64). Moreover, QS is crucial in both *V. cholerae* habitats. Thus, we sought to investigate whether oxygen modulates *V. cholerae* QS. First, we assessed the relative levels of the three known QS AIs from *V. cholerae* C6706 Sm^R^ (hereafter wild-type; WT) following aerobic and anaerobic growth. AI activity in cell-free culture fluids was measured using a set of three bioluminescent *V. cholerae* strains, each of which exclusively reports on one QS AI (either AI-2, CAI-1, or DPO) when it is supplied exogenously.

Unlike *V. cholerae* cultured in the presence of oxygen (hereafter +O_2_), *V. cholerae* grown in the absence of oxygen (hereafter -O_2_) produced no CAI-1. Twice as much AI-2 and DPO accumulated in *V. cholerae* cultured -O_2_ than +O_2_ (Figure 2A-C). We note that the dynamic ranges for the CAI-1 and DPO assay are ∼1,000- and ∼4-fold, respectively, while that for the AI-2 assay is ∼100,000-fold (39, 47). Thus, we consider the changes in CAI-1 and DPO to be physiologically relevant, whereas that for AI-2 is likely not, so we do not consider AI-2 further in this work. Additionally, *V. cholerae* cultured -O_2_ grew to a lower final cell density than when grown +O_2_ (Supplementary Figure 1A). We controlled for the reduced cell growth that occurs in the -O_2_ conditions, nonetheless, no CAI-1 could be detected (Supplementary Figure 1B). Beyond lacking O_2_, our culture medium lacked an alternative terminal electron acceptor. Thus, we also considered the possibility that *V. cholerae* cultured in -O_2_ conditions was unable to respire and therefore unable to drive CAI-1 generation. However, supplementation of the *V. cholerae* -O_2_ cultures with the alternative terminal electron acceptor fumarate, which is readily consumed by *V. cholerae* (4), did not rescue CAI-1 production (Supplementary Figure 1B). Collectively, these data suggest that production of CAI-1 and DPO by *V. cholerae* is affected by oxygen levels. In the remainder of this study, we focus on the functioning of the DPO-VqmA QS circuit under different conditions that are predicted to be encountered in the host. We address possible ramifications of our results concerning CAI-1 and AI-2 in the Discussion.

**Figure 2.**
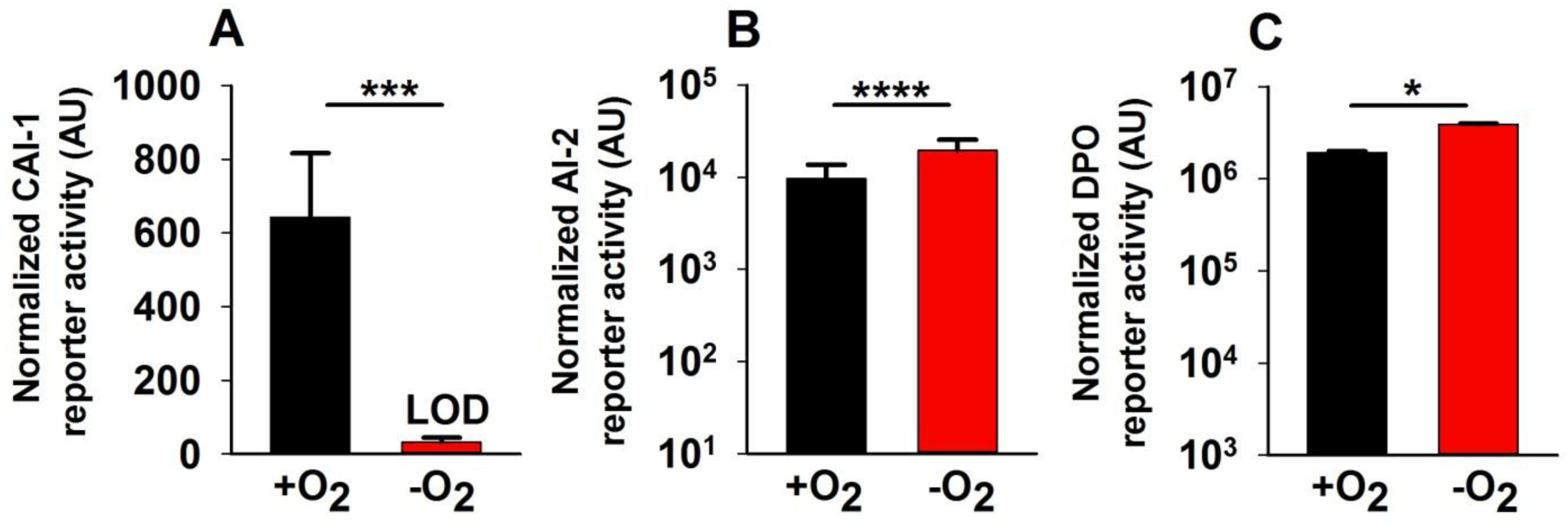
Oxygen deprivation modulates *V. cholerae* AI production. **(A)** 80%, **(B)** 25%, or **(C)** 30% cell-free culture fluids prepared from WT *V. cholerae* grown in the presence or absence of O_2_ were provided to *V. cholerae* reporter strains that produce bioluminescence in response to exogenous **(A)** CAI-1, **(B)** AI-2, or **(C)** DPO. Data represent the average values of biological replicates (*n*=3) and error bars represent SD. * denotes p<0.05, *** denotes p<0.001, and **** denotes p<0.0001.

### VqmA exhibits increased activity in the absence of oxygen

Given that *V. cholerae* accumulated more DPO under -O_2_ conditions than +O_2_ conditions, we wondered whether the VqmA-DPO QS system would, in turn, display increased activity under -O_2_ conditions compared to +O_2_ conditions. VqmA controls the expression of the *vqmR* gene, encoding the small RNA VqmR (Figure 1). Therefore, expression of a *vqmR*-*lacZ* transcriptional fusion can be used to assess VqmA activity (47). Beta-galactosidase was selected as the reporter because its activity is not affected by oxygen. The *vqmR-lacZ* construct was integrated onto the chromosome of Δ*tdh V. cholerae*. Tdh (threonine dehydrogenase) is required for DPO production (47). Thus, the Δ*tdh* strain makes no DPO but activates *vqmR-lacZ* expression in response to exogenously supplied DPO. We measured activity following growth in +O_2_ and -O_2_ conditions and in the absence and presence of exogenous DPO. In the presence of O_2_, *vqmR-lacZ* activity increased following supplementation with DPO (Supplementary Figure 2A). Compared to the +O_2_ conditions, *vqmR-lacZ* activity was higher under -O_2_ conditions, both in the absence and presence of DPO (Supplementary Figure 2A). Increased DPO-independent *vqmR-lacZ* expression in the absence of O_2_ could be a consequence of increased production of VqmA or increased VqmA activity. To distinguish between these possibilities, we first examined whether changes in O_2_ levels alter VqmA production by quantifying VqmA-FLAG produced from the chromosome - /+O_2_ and -/+ DPO. Similar levels of VqmA-FLAG were produced in all cases, suggesting that a change in VqmA abundance does not underlie increased *vqmR-lacZ* expression under -O_2_ conditions. (Supplementary Figure 2B). We next tested O_2_-driven changes in VqmA activity. To do this, we uncoupled expression of *vqmA-FLAG* from its native promoter by cloning *vqmA-FLAG* onto a plasmid under an arabinose inducible promoter (hereafter, p*vqmA-FLAG*). We introduced the plasmid into a Δ*vqmA* Δ*tdh V. cholerae* strain harboring the *vqmR-lacZ* chromosomal reporter and we measured both β- galactosidase output as well as VqmA-FLAG abundance in the same samples. VqmA- FLAG levels did not change under the different conditions (Figure 3A), however, *vqmR- lacZ* reporter activity normalized to cellular VqmA-FLAG levels increased in the cells exposed to DPO, and overall activity was ∼4-7-fold higher under -O_2_ conditions than +O_2_ conditions both in the presence and absence of DPO (Figure 3B). We conclude that VqmA displays an increased capacity to activate gene expression under anaerobic conditions relative to aerobic conditions.

**Figure 3.**
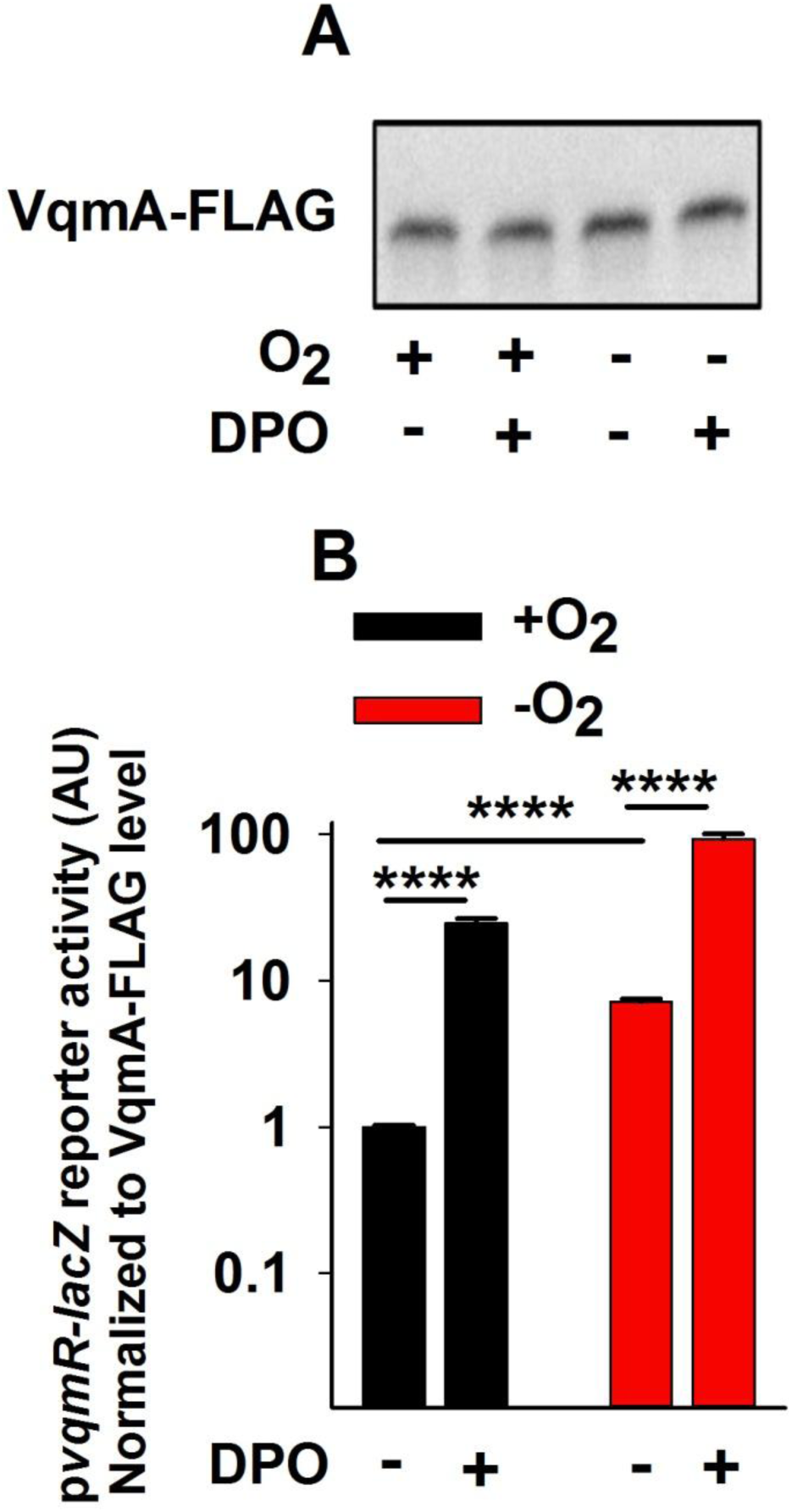
Apo- and Holo-VqmA activities are enhanced in the absence of oxygen. **(A)** Western blot showing VqmA-FLAG abundance in the Δ*vqmA* Δ*tdh V. cholerae* carrying p*vqmA-FLAG* and treated as shown for 1 h. DPO was supplied at 25 µM DPO. **(B)** Transcriptional activity for the p*vqmR-lacZ* reporter from the same samples presented in A. Data in B were normalized to the level of VqmA-FLAG in A. In B, data represent the average values of biological replicates (*n*=3) and error bars represent SD. **** denotes p<0.0001.

### VqmA forms intra- and inter-molecular disulfide bonds in an oxygen- and DPO- dependent manner

We wondered what molecular mechanism drives the increase in VqmA activity under -O_2_ conditions (Figure 3). The cytoplasmic compartment of aerobically respiring *V. cholerae* is relatively oxidizing (35, 40). Thus, decreased oxygen levels would shift the cytoplasm to a reducing environment (42). Proteins can respond to such changes via redox-responsive cysteine residues (35, 38). Inspection of the VqmA amino acid sequence revealed the presence of four cysteine residues and all are strictly conserved in VqmA homologs in other vibrio species, but not in the VqmA receptor recently discovered in a vibriophage (Supplementary Figure 3). These findings led us to consider a model in which, in addition to activation by DPO, VqmA activity is regulated by redox-responsive cysteine residues.

Cysteine residues often undergo disulfide bond formation (1, 38, 48). We assessed whether VqmA forms disulfide bonds *in vivo* and, if so, whether their formation is influenced by DPO and/or oxygen levels. We grew Δ*vqmA* Δ*tdh V. cholerae* carrying the p*vqmA-*FLAG construct under +O_2_ conditions. We subsequently divided the culture into four aliquots. One portion was untreated (+O_2_, -DPO), one portion was supplied DPO (+O_2_, +DPO), one portion was deprived of oxygen (-O_2_, -DPO), and one portion was deprived of oxygen and supplemented with DPO (-O_2_, +DPO). We extracted protein and analyzed the VqmA-FLAG protein profiles by immunoblot. These analyses were performed with or without the addition of the reductant β- mercaptoethanol (BME) to distinguish between VqmA-FLAG species that had and had not formed disulfide bonds. Previous studies have shown that the presence of an intra-molecular disulfide bond leads to a protein species displaying increased gel mobility compared to the same protein lacking the bond (2, 55). By contrast, inter-molecular disulfide bonds produce cross-linked protein oligomers that migrate with slower mobility than the corresponding monomers (16, 44). We first consider the results for VqmA under +O_2_ conditions: Under non-reducing conditions (-BME) and in the absence of DPO, VqmA-FLAG displayed mobility consistent with an oxidized monomer (labeled O- Monomer; Figure 4A and 4B lane 1). Treatment with BME caused VqmA-FLAG to migrate more slowly, consistent with it being a reduced monomer (labeled R-Monomer; Figure 4A and 4B lane 2). Administration of DPO drove formation of an additional VqmA-FLAG species, corresponding in size to an oxidized dimer (labeled O-Dimer, Figure 4A and 4B lane 3), but only under oxidizing (i.e., -BME) conditions (Figure 4B, compare lanes 3 and 4). These results suggest that, under aerobic conditions, a fraction of VqmA harbors an intra-molecular disulfide bond and DPO-bound VqmA forms an inter-molecular disulfide bond. Under anaerobic conditions, the portion of VqmA containing the intra-molecular disulfide bond decreased (Figure 4B compare lane 1 to lane 5 and lane 5 to lane 6) while the DPO-dependent inter-molecular disulfide bonded species was unaffected by the absence of oxygen (Figure 4A compare lane 3 to lane 7 and lane 7 to lane 8).

**Figure 4.**
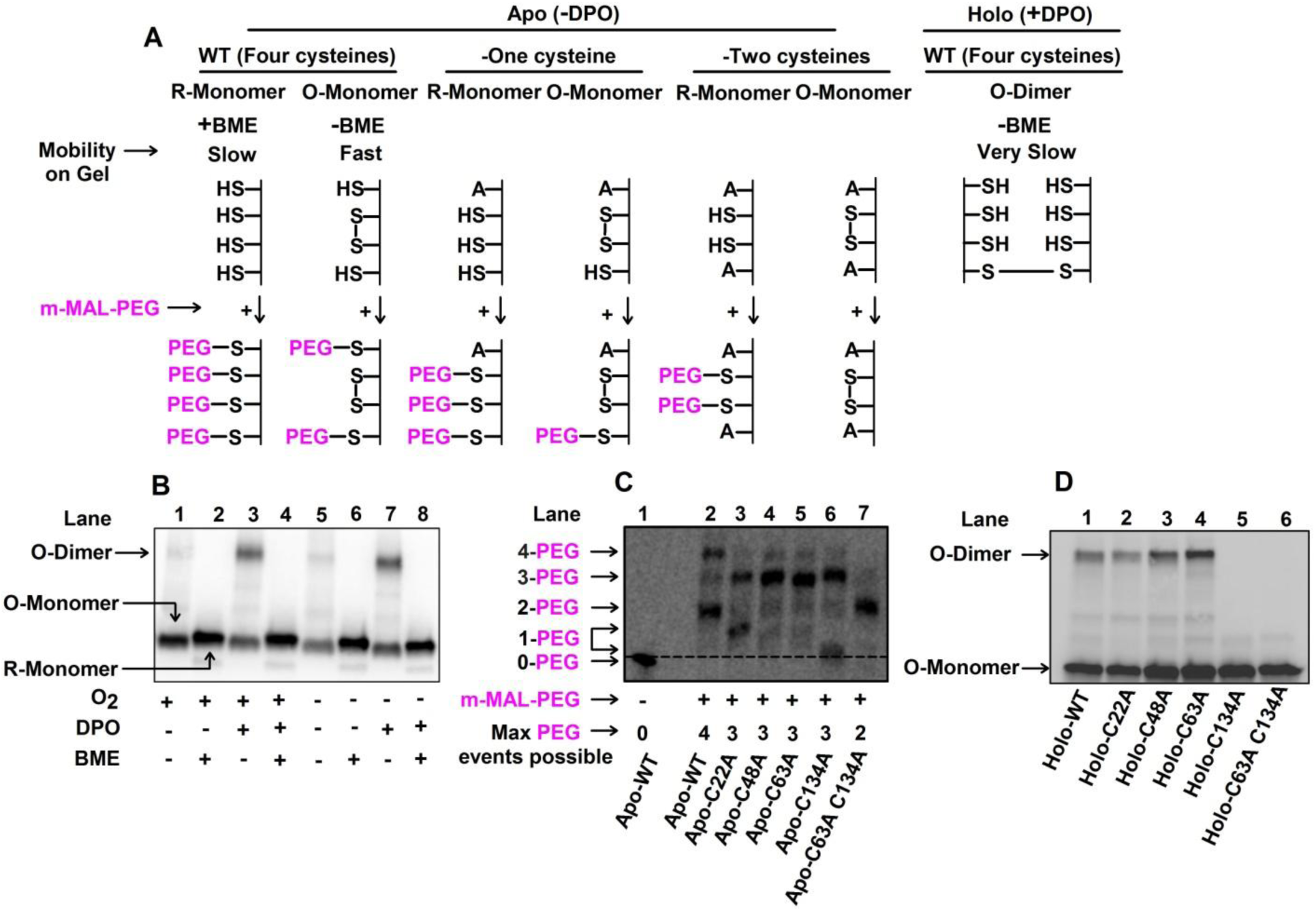
VqmA forms intra- and inter-molecular disulfide bonds in an oxygen- and DPO-dependent manner. **(A)** Schematic depicting the oxidized and reduced forms of VqmA (top) and the strategy employed to interrogate intra-molecular disulfide bond formation by trapping reduced thiols with methoxypolyethylene glycol maleimide (m- MAL-PEG; 5kDa, bottom) (32). **(B)** Western blot showing VqmA-FLAG protein produced by the Δ*vqmA* Δ*tdh V. cholerae* strain carrying p*vqmA-FLAG* following the specified treatments. **(C)** Western blot showing VqmA-FLAG protein produced by the Δ*vqmA* Δ*tdh V. cholerae* strain carrying the designated p*vqmA-*FLAG alleles following treatment with m-MAL-PEG (32) in the absence of DPO. Note that in lanes 3 and 6, the proteins containing 1-PEG modification migrate somewhat differently. We infer the bands to be 1-PEG events by comparison with the 0-PEG event in lane 1 and the 2-PEG event in lane 7. A likely explanation is that the protein containing 1-PEG in lane 6 adopts a more compact conformation than the modified protein in lane 3 and so it migrates faster through the gel (2, 55). The dotted line distinguishes the 0-PEG VqmA from the lowest 1-PEG decorated species. **(D)** Western blot showing VqmA-FLAG for the strains in C following supplementation with 25 µM DPO under non-reducing conditions.

To garner additional evidence for the presence of an intra-molecular disulfide bond(s) in the species designated VqmA O-monomer in Figure 4B, we treated samples prepared from cells grown under +O_2_ and -DPO conditions with methoxypolyethylene glycol maleimide (m-MAL-PEG) (32). m-MAL-PEG alkylates reduced cysteine residues and in so doing, confers an ∼5 kDa molecular weight change for each alkylation event (see schematic in Figure 4A). VqmA contains four cysteine residues, thus the fully reduced protein would undergo 4 m-MAL-PEG events while if one intra-molecular disulfide bond exists, only two cysteine residues could react. Following treatment, Apo-VqmA-FLAG migrated primarily as two bands, corresponding to four (∼40% of protein) and two (∼60% of protein) alkylation events (Figure 4C, compare lane 1 to lane 2), confirming that while a portion of VqmA-FLAG is fully reduced *in vivo*, the majority of the protein exists as an oxidized species containing one intra-molecular disulfide bond.

To determine the residues involved in VqmA disulfide linkages, we conducted two experiments. First, we employed an *in vitro* thiol-trapping strategy based on sequential reactions that modify accessible cysteine residues with two thiol-specific reagents (Supplementary Figure 4A) (52). In the initial thiol-blocking step, purified 6His- VqmA treated with diamide, a thiol-specific oxidant (30), was denatured and incubated with chloroacetamide (CAA). CAA alkylates accessible cysteine residues (i.e., those not involved in disulfide bonds), blocking them from further modification. The CAA treated sample was next treated with TCEP, a reductant, enabling non-CAA labeled cysteines to be reduced. These newly freed residues were labeled in the final alkylation step with N-ethylmaleimide (NEM). The sample was then analyzed by mass spectrometry. The logic is that if a particular cysteine residue was inaccessible to CAA due to disulfide bonding, it would be preferentially labeled with NEM in the subsequent NEM modification step. Thus, the NEM/CAA ratio would be >1. By contrast if a cysteine residue was not involved in disulfide modification, it would be preferentially labeled with CAA and have a NEM/CAA ratio < 1. The VqmA C48 and C63 residues had NEM/CAA ratios of ∼100 and ∼10, respectively, suggesting both of these residues are involved in disulfide-linkages (Supplementary Figure 4B). We were unable to obtain good coverage of the C134 and C22 residues using this technique so we could not similarly assess them.

Our second experiment to probe disulfide bonds in VqmA relied on mutagenesis. We individually substituted an alanine residue for each cysteine residue in the p*vqmA- FLAG* construct, introduced the plasmids into the Δ*vqmA* Δ*tdh V. cholerae* strain, and repeated the analyses described in Figure 4C. We first consider the case of intra-molecular disulfide bond formation under +O_2_ -DPO conditions. In the mutant proteins, following replacement of a cysteine residue with alanine, a maximum of three residues can react with m-MAL-PEG in the fully reduced protein (Figure 4A, top). However, if an intra-molecular disulfide bond is present, then only one cysteine residue can be decorated with m-MAL-PEG. Figure 4C shows that the Apo-VqmA C22A and Apo-VqmA C134A proteins each migrated as two bands, a result consistent with portions of each protein harboring one and three m-MAL-PEG moieties (Figure 4C; compare lane 3 to lanes 2 and 1; and lane 6 to lanes 2 and 1). This result suggests that the fraction of Apo-VqmA C22A and the fraction of Apo-VqmA C134A that exhibit one m-MAL-PEG decoration harbor intra-molecular disulfide bonds. Apo-VqmA C48A and Apo-VqmA C63A migrated largely as single bands at the region corresponding to three m-MAL-PEG decorations (Figure 4C; compare lanes 4 and 5 to lane 2), suggesting that these proteins exist as reduced species and are thus incapable of forming intra-molecular disulfide bonds. Therefore, we conclude that in WT VqmA, an intra-molecular disulfide bond is formed between cysteine residues 48-63.

Next, we consider inter-molecular disulfide bond formation under +O_2_ +DPO conditions, and under non-reducing conditions (i.e., -BME). Figure 4D shows that the Holo-VqmA C22A, Holo-VqmA C48A, and Holo-VqmA C63A proteins migrated as mixtures of oxidized monomers and oxidized dimers, while the Holo-VqmA C134A protein migrated exclusively as an oxidized monomer (compare lanes 2-5 to lane 1). These data suggest that in Holo-VqmA, there is a C134-C134 inter-molecular disulfide linkage. We also constructed and assessed the double VqmA C63A C134A mutant under +O_2_ +m-MAL-PEG and +O_2_ +DPO conditions. Figure 4C shows that under aerobic conditions, all of the Apo-VqmA C63A C134A protein contains two m-MAL-PEG decorations (compare lane 7 to lane 2), confirming that the two remaining cysteine residues were accessible and that the protein is fully reduced. Figure 4D shows that under aerobic conditions Holo-VqmA C63A C134A migrates entirely as a monomer (compare lanes 6 to lane 1). Thus, Holo-VqmA C63A C134A is incapable of forming both intra- and inter-molecular disulfide bonds.

Collectively our data suggest that: 1) VqmA forms disulfide bonds *in vivo* and *in vitro*; 2) an intra-molecular disulfide bond is formed between VqmA C48-C63, and an inter-molecular disulfide bond is made between C134-C134, and 3) disulfide bond formation is influenced by both oxygen and DPO.

### VqmA activity is limited by the intra-molecular disulfide bond and enhanced by the inter-molecular disulfide bond

To explore the *in vivo* consequences of VqmA disulfide bond formation on VqmA function, we tested whether the Apo- and Holo- mutant VqmA proteins that are incapable of forming particular intra- and/or inter-molecular disulfide bonds displayed altered abilities to activate target *vqmR* transcription (see schematic in Supplementary Figure 5A). We introduced the *vqmA- FLAG, vqmA C48A-FLAG* and the *vqmA C134A-FLAG* alleles onto the chromosome of a *V. cholerae* Δ*tdh* strain carrying *vqmR-lacZ* and measured reporter activity following aerobic growth +/- DPO. Our rationale was that WT VqmA forms both the C48-C63 intra-molecular bond and the C134-C134 inter-molecular bond. By contrast the VqmA C48A variant is unable to form the C48-C63 intra-molecular bond and the VqmA C134A variant is unable to form the C134-C134 inter-molecular bond (from Figure 4C and 4D). Therefore, by comparing the activities of these three proteins, we could assess the effect of individually eliminating each disulfide bond on VqmA activity. We likewise made a strain carrying *vqmA C63A C134A-FLAG* on the chromosome to examine the effect of simultaneous elimination of both disulfide bonds.

**Figure 5.**
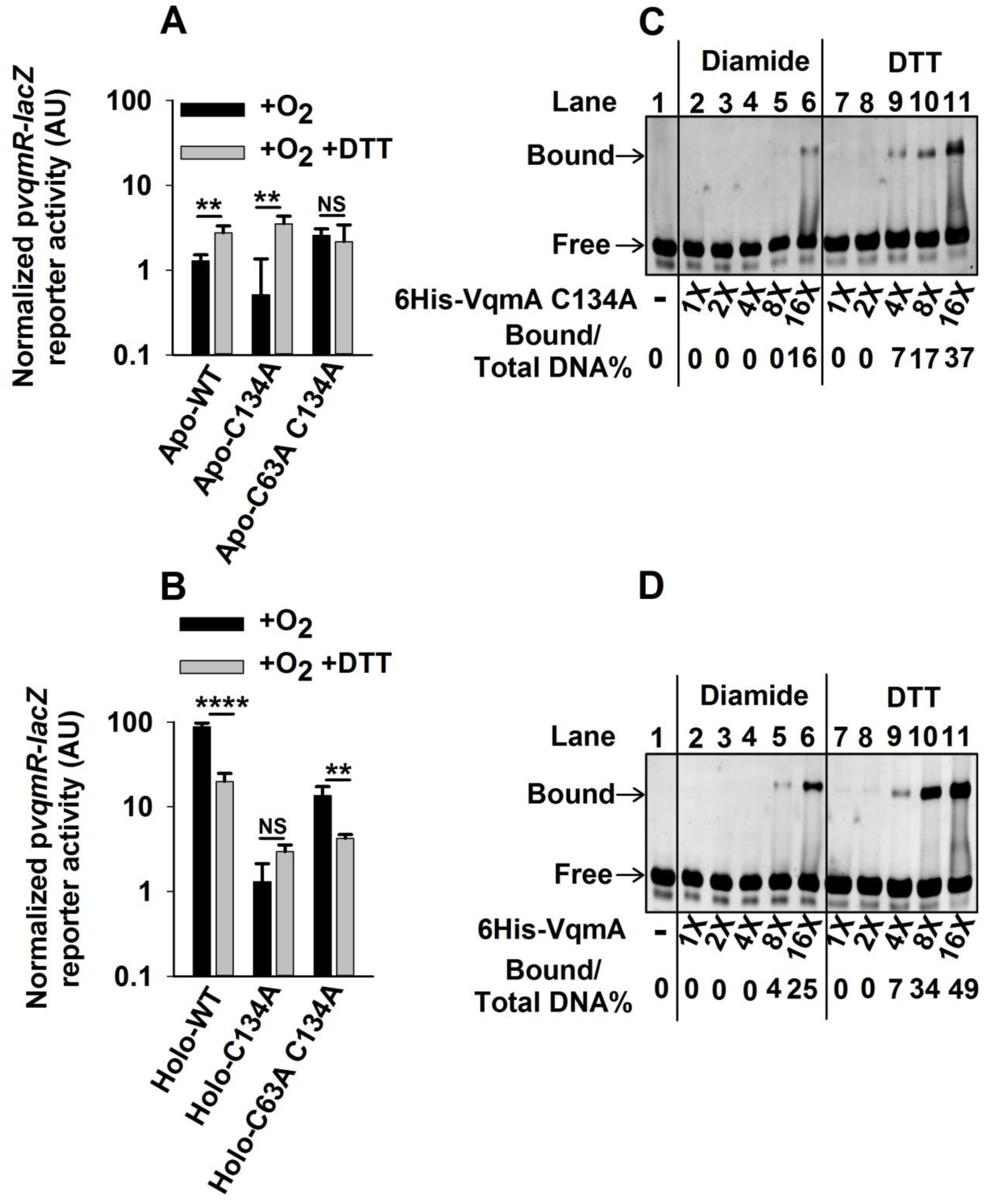
VqmA activity is differentially modulated by the cellular redox environment and intra- and inter-molecular disulfide bonds. **(A)** p*vqmR-lacZ* activity in Δ*vqmA* Δ*tdh V. cholerae* carrying the designated p*vqmA-FLAG* constructs following growth in the presence of O_2_ without (black) or with DTT (gray). **(B)** Strains cultured as in A supplemented with 25 µM DPO. **(C)** Electromobility shift analysis (EMSA) of 6His-VqmA C134A binding to *vqmR* promoter DNA. **(D)** EMSA as in C for WT 6His-VqmA. In C and D, all lanes contained 35 ng of promoter DNA and, where indicated, dilutions of protein were used with 16X = 2 µg. Bound and free correspond to DNA that is and is not bound to VqmA protein, respectively. As indicated, VqmA had been treated with 10-fold molar excess of diamide or DTT. Data in A and B represent average values of biological replicates (*n*=3) and error bars represent SD. ** denotes p<0.01, **** denotes p<0.0001, and NS denotes p>0.05.

First we consider the case of Apo-VqmA. Apo-VqmA C48A and Apo-VqmA C134A exhibited an ∼2-fold and ∼4-fold increase and decrease, respectively, in reporter activity, relative to WT Apo-VqmA (Supplementary Figure S5B). Apo-VqmA C63A C134A exhibited a ∼3-fold increase in reporter activity relative to WT Apo-VqmA and a ∼10-fold increase in reporter activity relative to the strain carrying the Apo-VqmA C134A single mutant (Supplementary Figure S5B). Thus, we conclude that the C48-C63 intra-molecular disulfide bond limits transcriptional activity of Apo-VqmA.

Next, we consider the case for Holo-VqmA. In cultures supplemented with DPO, *vqmR-lacZ* reporter activity increased ∼20-30-fold, in a DPO-concentration-dependent manner in the strain carrying WT Holo-VqmA relative to the strain with WT Apo-VqmA (Supplementary Figure S5C). The strain carrying the Holo-VqmA C48A variant displayed a further increase in DPO-dependent reporter activity relative to the WT Holo- VqmA. However, in strains harboring Holo-VqmA C134A and Holo-VqmA C63A C134A, only modest (3-4-fold) responses to DPO occurred (Supplementary Figure S5C). Thus, the DPO-responsive, C134-C134 inter-molecular disulfide-bond enhances VqmA transcriptional activation activity.

### VqmA activity is differentially modulated by the cellular redox environment

To test whether VqmA activity is responsive to cellular redox, we supplemented the strains carrying the different VqmA variants with DTT, a cell-permeable reductant. We reasoned that if the absence of oxygen generates a reducing environment that prevents the formation of a particular disulfide bond, addition of DTT would mimic this condition by promoting a reducing cellular environment, including in the presence of oxygen. To control for any potential DTT-induced changes in the levels of chromosomally expressed *vqmA*, we introduced plasmids harboring arabinose inducible *vqmA-FLAG, vqmA C134A*-*FLAG*, or *vqmA C63A C134A-FLAG* into the *V. cholerae* Δ*vqmA* Δ*tdh* strain carrying *vqmR-lacZ* and measured reporter activity +/- DPO and +/- DTT.

In the absence of DTT, the Apo- and Holo- plasmid-borne VqmA variants displayed reporter activities similar to when the variants were expressed from the chromosome (plasmid-borne variants are in Figure 5A and 5B, compare the black bars in each panel; for chromosomal variants, see Figure S5B and C). Treatment with DTT increased reporter activity for WT Apo-VqmA and Apo-VqmA C134A but did not alter the reporter activity in the strain carrying Apo-VqmA C63A C134A (Figure 5A, compare black and gray bars for each strain). As a reminder, WT Apo-VqmA and Apo-VqmA C134A form the C48-C63 intra-molecular disulfide bond, while the Apo-VqmA C63A C134A protein does not. Thus, these data suggest that a reducing environment interferes with formation of the C48-C63 intra-molecular disulfide bond, thereby eliminating its negative effect on Apo-VqmA activity.

Regarding WT Holo-VqmA, reporter activity diminished by ∼6-fold when DTT was present in addition to DPO (Figure 5B; compare the first pair of black and gray bars). In contrast, DTT supplementation did not significantly affect reporter activity in the strain carrying Holo-VqmA C134A (Figure 5B; compare the second set of black and gray bars), while ∼3-fold lower activity was produced by the strain carrying Holo-VqmA C63A C134A (Figure 5B; compare the third set of black and gray bars). Since the activity of VqmA C134A was not affected by DTT supplementation, we conclude that the C134- C134 inter-molecular bond does not form in a reducing environment, and without that bond, Holo-VqmA transcriptional activity is diminished.

### VqmA DNA binding capacity is differentially modulated by its redox environment

We suspected that the *in vivo* redox-dependent changes in VqmA disulfide bond formation would have ramifications on VqmA DNA binding capability. To explore this notion, we examined the ability of purified VqmA proteins to bind p*vqmR* promoter DNA. First, to examine the role of the C48-C63 intra-molecular disulfide bond, we assessed DNA binding for 6His-VqmA C134A treated with diamide (to enable disulfide bond formation) or DTT (to prevent disulfide bond formation) (see Supplementary Figure 6). Relative to the DTT treated protein, the diamide treated protein exhibited a 4-fold reduction in DNA binding (Figure 5C; compare lanes 2-5 with lanes 7-10). This result suggests that formation of the C48-C63 intra-molecular disulfide bond limits DNA binding.

**Figure 6.**
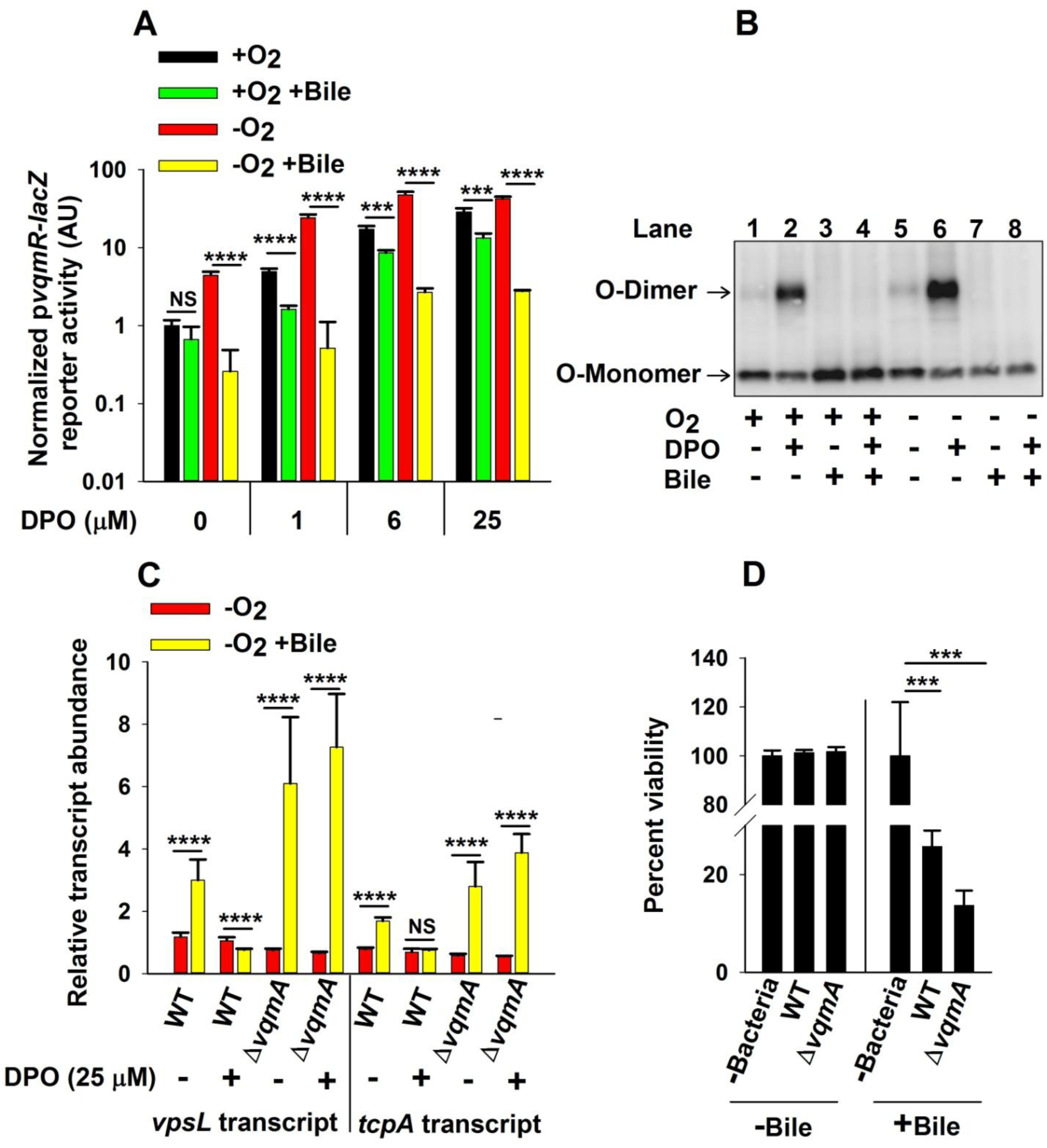
Bile salts disrupt VqmA-DPO-mediated signal transduction and promote *V. cholerae* virulence. **(A)** p*vqmR-lacZ* activity in Δ*vqmA* Δ*tdh V. cholerae* carrying p*vqmA- FLAG* following 1 h in the presence or absence of O_2_, 0.5% (v/v) bile, and 1-25 µM DPO or combinations of all three treatments as indicated. **(B)** Western blot showing VqmA- FLAG for the strain in A following the indicated treatments. **(C)** Relative expression levels of *vpsL* and *tcpA* in WT and Δ*vqmA V. cholerae* following the designated treatments. **(D)** Viability of human intestinal CaCo-2 cells in the absence of bacteria or following challenge by WT or Δ*vqmA V. cholerae* and in the presence or absence of 0.05% bile. Data in panels A, and D represent average values of biological replicates (*n*=3) and error bars represent SD. Data in panel C represent the average value of three biological replicates and two technical replicates for each sample (*n*=6) and error bars represent SD. *** denotes p<0.001, **** denotes p<0.0001, and NS denotes p>0.05.

We investigated the role of the VqmA C134-C134 inter-molecular disulfide bond on DNA binding by comparing the DNA binding capabilities of 6His-VqmA and 6His- VqmA C134A. Diamide treated 6His-VqmA was approximately twice as potent at binding DNA as was 6His-VqmA C134A [compare left halves of Figure 5C (VqmA C134A) and Figure 5D (WT VqmA)], suggesting that the C134-C134 inter-molecular disulfide bond promotes DNA binding. The DTT treated 6His-VqmA C134A and 6His- VqmA proteins showed no difference in DNA binding capability [compare right halves of Figure 5C (VqmA C134A) and Figure 5D (WT VqmA)]. Like 6His-VqmA C134A, DTT treated 6His-VqmA was more proficient in DNA binding than diamide treated 6His- VqmA, consistent with intra-molecular disulfide bond formation limiting DNA binding (Figure 5D). Collectively, the data in Figures 4 and 5 suggest a model in which the transcriptional and DNA binding activities of both Apo-VqmA and Holo-VqmA are modulated by disulfide bond formation, the cytoplasmic redox environment, and by the level of O_2_ in the environment.

### Bile salts interfere with VqmA disulfide-bond formation and decrease VqmA transcriptional activation activity

Our above findings suggest that the VqmA-DPO signal transduction pathway, which represses virulence factor production and biofilm formation, is most highly active under anaerobic conditions. *V. cholerae* encounters anaerobiosis in the human intestine. The paradox is that in the intestine, *V. cholerae* is virulent and makes biofilms. We thus wondered if a possible host intestinal signal(s) could modulate VqmA-DPO signaling, allowing infection to proceed under anaerobic conditions. Bile salts, present in high concentrations in the human small intestine, can alter the redox environment of bacterial cells and thereby affect disulfide bond formation in cytoplasmic proteins (11). Thus, we were inspired to investigate whether bile salts could abrogate VqmA-DPO transcriptional activation activity. We cultured the Δ*vqmA* Δ*tdh V. cholerae* strain carrying the arabinose inducible *pvqmA-FLAG* construct and *vqmR-lacZ* on the chromosome in the presence and absence of oxygen, bile, and DPO and we measured reporter activity. Bile treatment caused ∼2- fold and ∼10-fold decreases in *vqmR-lacZ* reporter activity under +O_2_ and -O_2_ growth, respectively (Figure 6A; first four bars). Bile supplementation also decreased *vqmR-lacZ* reporter activity in cultures supplied with DPO, again with the maximum effect observed under - O_2_ growth (Figure 6A; 5^th^ bar onward). To test whether the presence of bile affects VqmA disulfide bonds, we used analyses similar to those in Figure 4B. Consistent with the *vqmR-lacZ* reporter activity, bile supplementation prevented formation of O-dimers, both in the presence and absence of O_2_, suggesting it interferes with the Holo-VqmA C134-C134 inter-molecular disulfide bond (Figure 6B).

### Bile salt-mediated disruption of VqmA-DPO-driven signal transduction promotes *V. cholerae* virulence

In *V. cholerae*, VqmA-DPO-directed production of VqmR results in decreased expression of genes involved in biofilm formation and virulence factor expression, including *vpsL* and *tcpA*, respectively (46, 47). VpsL is required to synthesize *V. cholerae* exopolysaccharide, an essential component of the biofilm matrix, and TcpA is a virulence factor required for *V. cholerae* to colonize the human small intestine (17, 57). Our finding that bile supplementation inhibited VqmA-DPO function and that the effect of bile-mediated inhibition occurred primarily in the absence of oxygen, led us to predict that the repression of *vpsL* and *tcpA* expression would also be maximally disrupted following bile supplementation in the absence of oxygen. We measured transcript levels of *vpsL* and *tcpA* in the WT and Δ*vqmA* strains following exposure to 25 µM DPO, bile, deprivation of oxygen, or combinations of the three treatments.

In WT *V. cholerae*, bile supplementation modestly increased *vpsL* and *tcpA* transcript levels (∼3- and ∼2-fold, respectively) however, only under -O_2_ -DPO conditions (Supplementary Figure 7 and Figure 6C). These data suggest that bile induces genes required for biofilm formation and virulence in the absence of oxygen and Holo-VqmA can override the effect of bile. Transcript levels for both *vpsL* and *tcpA*, under -O_2_ -DPO conditions were further increased in the Δ*vqmA* strain treated with bile (∼6- and ∼3-fold, respectively). We interpret this result to mean that bile also induces an increase in *vpsL* and *tcpA* expression through a pathway that does not involve VqmA. However, because the major effect of bile occurs only in the Δ*vqmA* strain, we conclude that this additional pathway is epistatic to VqmA in the control of *vpsL* and *tcpA*.

**Figure 7.**
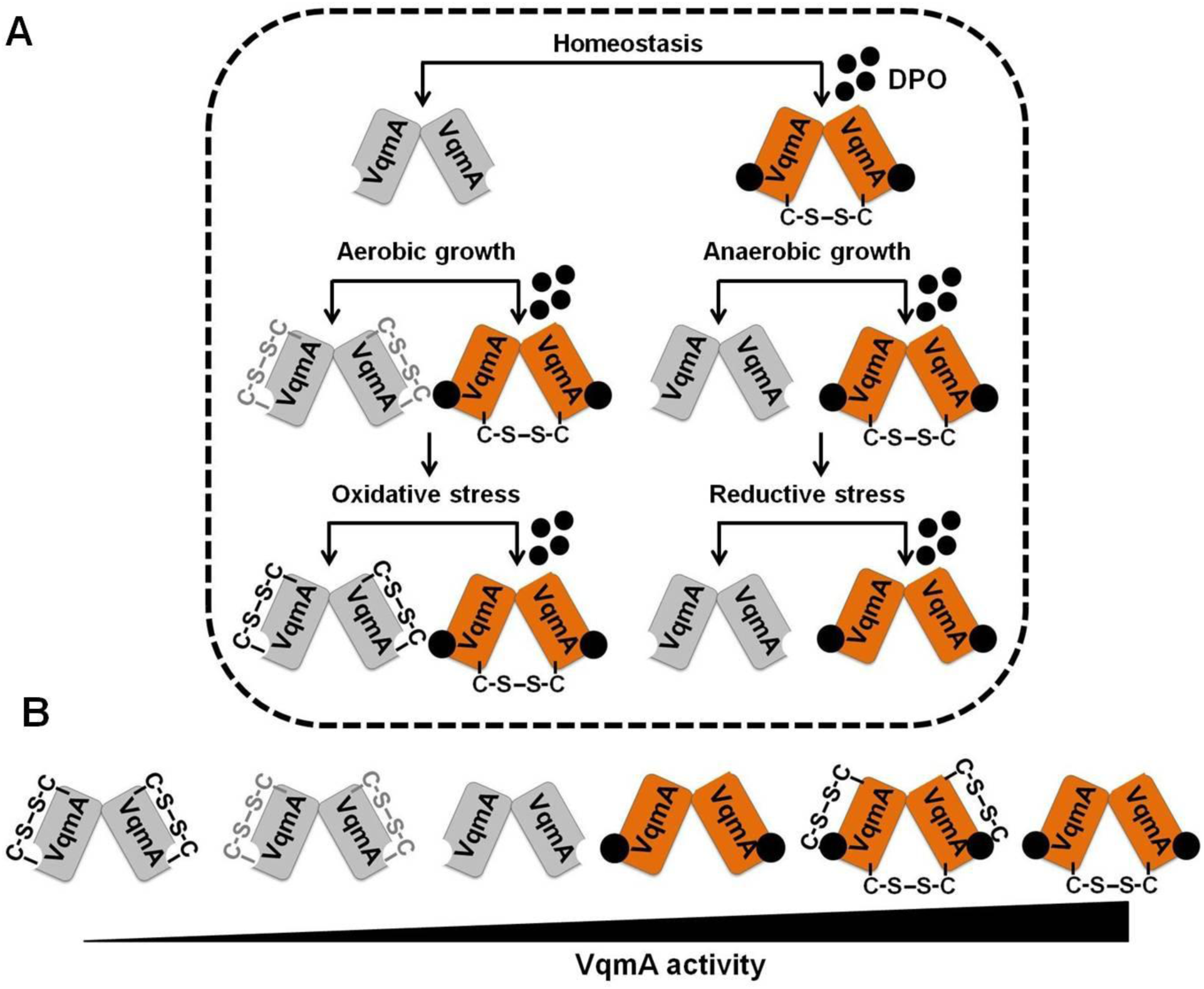
Model depicting VqmA as a hub protein that compiles quorum-sensing, environmental, and host information. **(A)** VqmA can exist in different states *in vivo* depending upon the availability of the DPO ligand and the cellular redox state. Thus, Apo-VqmA forms the C48-C63 intra-molecular disulfide bond that suppresses its ability to bind DNA. Apo-VqmA activity is high under reducing growth conditions in which formation of the intra-molecular C48-C63 bond is inhibited. Holo-VqmA forms the C134- C134 inter-molecular disulfide bond that promotes DNA binding. Reductive stress disrupts the formation of the inter-molecular disulfide bond. We propose that growth in the presence of bile salts imposes reductive stress, disrupts the formation of the C134- C134 inter-molecular disulfide bond, and restricts VqmA DNA binding, thereby promoting virulence and biofilm formation. The gray and black intra-molecular disulfide bonds denote partially-oxidized and fully-oxidized VqmA, respectfully. **(B)** Relative VqmA activity levels as a consequence of disulfide bond formation. For simplicity, the fourth and fifth species in B are not displayed in A.

We tested whether the above changes in gene expression translated into alterations in *V. cholerae* pathogenicity by assaying whether bile affected cytotoxicity of *V. cholerae* in co-culture with human Caco-2 intestinal cells. We generated differentiated monolayers of Caco-2 cells and co-cultured them with either WT or Δ*vqmA V. cholerae* in the presence and absence of bile. The presence of bile, increased *V. cholerae*-mediated cytotoxicity to Caco-2 cells with the Δ*vqmA* strain driving twice as much killing as WT *V. cholerae* (Figure 6D). Collectively, our data suggest that bile salts induce increased *V. cholerae* virulence when bacteria are deprived of oxygen, part of the bile effect is exerted via interference with the VqmA-DPO-VqmR QS circuit, and the presence of VqmA limits the ability of bile to affect target gene expression.

## Discussion

The diversity of environments that *V. cholerae* inhabits, from the ocean to marine organisms to the human stomach to the human intestine, necessitates that the bacterium rapidly perceives changes in its external environment and appropriately tailors its gene expression programs. Our current study reveals that *V. cholerae* alters both which AIs are produced and the functioning of the VqmA-DPO-VqmR QS circuit in response to its environment. To our knowledge, these findings represent the first dissection of QS activity in the absence of oxygen in a facultative aerobic bacterium. We find that 1) the amount of an AI (CAI-1) produced is dictated by oxygen levels; 2) a single QS protein (VqmA) is capable of integrating information from three sources (AI, oxygen, bile); and 3) two disulfide bonds in a QS receptor (VqmA) antagonize one another with respect to their effects on protein activity, thereby aiding in perception and response to changes in cellular redox.

The benefit(s) of producing and detecting multiple QS AIs has long been mysterious with respect to *V. cholerae* biology. Evidence suggests that each AI conveys specific information into the cell: CAI-1 measures the abundance of vibrios (kin) and AI- 2 and DPO measure the level of non-vibrio (non-kin) in the vicinity (5, 46, 47). Our finding that CAI-1 is not produced in the absence of oxygen suggests that CAI-1 may also convey information about the external environment. Strains lacking the ability to synthesize CAI-1 display reduced survival in seawater and following challenge with oxidative stress (25). Thus, we propose that CAI-1 could drive the expression of genes required for the aerobic segment of the *V. cholerae* lifecycle and we are now testing this idea. The fact that CAI-1 is not produced under anoxia suggests that *V. cholerae* cannot take a census of kin in the absence of oxygen. Either kin counting is dispensable under anoxia or, perhaps, another molecule(s)/mechanism performs this function. By contrast, at a minimum based on sequencing data, thousands of bacterial species, including those found in the human microbiota, can synthesize AI-2, the AI used for inter-species communication (49). We found that AI-2 is synthesized in the absence of oxygen. Perhaps measurement of the abundance of non-kin bacteria is of paramount importance in densely populated niches containing complex bacterial consortia. Biosynthesis of both AI-2 and CAI-1 requires *S*-adenosylmethionine (SAM), an abundant metabolite that is crucial for methylation reactions (8, 28, 54). Thus, another possibility is that in *V. cholerae*, during periods of SAM limitation, CAI-1 production is curbed as a means of sparing SAM for other uses. Continued AI-2 production could suffice for QS-mediated cell-density tracking. Moreover, when SAM is used to produce AI-2, but not CAI-1, SAM is regenerated via downstream reactions (54, 61). Thus, making AI-2 from SAM would not deplete the SAM reservoir.

Oxygen is a terminal electron acceptor and therefore a critical substrate for bacterial growth. The human intestine is devoid of oxygen, and invading bacteria, such as *V. cholerae* that normally inhabit the relatively oxygenated marine environment, need to alter their physiology to survive. With the exception of a handful of studies (31, 33, 35), the molecular mechanisms by which *V. cholerae* perceives the absence of oxygen, and translates this information into changes in gene expression, are unexplored. Here, we demonstrate that, in the presence of O_2_, Apo-VqmA forms a C48-C63 intra-molecular disulfide bond that restricts the ability of the protein to bind DNA. Formation of this bond is inhibited in the absence of oxygen or following supplementation of aerobic cells with a reductant, that, analogous to the absence of oxygen, generates a reducing environment. Thus, we propose that by interacting with the cell’s redox environment, VqmA provides *V. cholerae* a mechanism to monitor oxygen levels (Figure 7A). In this context, we note that anaerobiosis causes a ∼7-fold increase in Apo-VqmA-dependent p*vqmR-lacZ* reporter activity (Figure 3B). By contrast, DTT supplementation under aerobic conditions causes only an ∼3-fold increase in Apo-VqmA activity while the activity of Apo-VqmA C63A C134A, that lacks both disulfide bonds, is unchanged (Figure 5A). One interpretation of these data is that Apo-VqmA is responsive to additional oxygen-dependent stimuli that are not mimicked by DTT or by the inability to make disulfide bonds. We are currently testing this possibility.

With respect to DPO-bound VqmA we found that Holo-VqmA forms a C134-C134 inter-molecular disulfide bond that promotes its ability to activate transcription. Formation of this bond is not modulated by oxygen levels, but is inhibited by the reductant DTT. The absence of oxygen imposes a mildly reducing environment on cells, while the presence of DTT imposes reductive stress (42, 58), suggesting that the C134- C134 inter-molecular VqmA disulfide bond may allow *V. cholerae* to monitor reductive stress. Collectively, we suggest a model in which cycling between multiple redox states, namely Oxidized-Apo-VqmA, Reduced-Apo-VqmA, Oxidized-Holo-VqmA, and Reduced-Holo-VqmA, enables *V. cholerae* to tune its QS-controlled collective behaviors to a range of redox states (Figure 7B). There exist examples of individual disulfide bonds restricting or enhancing the activity of transcription factors (7, 10, 65). To our knowledge, however, this is the first example in which the same protein simultaneously uses two different disulfide bonds to modulate activity.

Bile is an abundant compound in the human small intestine that is well known to alter virulence in *V. cholerae* and other enteric pathogens such as *Salmonella typhimurium* and *Shigella flexneri* (19, 43). Bile is a heterogeneous mixture of molecules and studies have largely focused on defining the roles of individual components in bacterial physiology. Intriguingly, the individual components can drive opposing effects. In *V. cholerae*, bile fatty acids repress while the bile salt taurocholate induces virulence (6, 9, 23, 36, 50, 63). In our current study, we elected to use a mixture of bile salts reasoning that this strategy would more closely approximate what *V. cholerae* encounters *in vivo*. Our data suggest that bile salts disrupt the formation of the VqmA C134-C134 inter-molecular disulfide bond. We do not know the mechanism by which this occurs. However, previous studies show that bile salts, specifically cholic acid (CHO) and deoxycholic acid (DOC), interfere with redox homeostasis in *Escherichia coli* by shifting the cellular environment to an oxidizing one and fostering disulfide bond formation in cytosolic proteins (11). In the context of our work, since VqmA inter-molecular disulfide bond formation is disrupted, we propose that application of a bile salts mixture to *V. cholerae* causes reductive stress. Consistent with this idea, taurocholate binds to and inhibits DsbA, a protein required for the introduction of disulfide bonds in periplasmic proteins (62). We currently do not know whether incubation of *V. cholerae* with CHO and DOC, rather than a bile salts mixture, would drive phenotypes mimicking those observed in *E. coli*.

What advantage does *V. cholerae* accrue by using the regulatory program uncovered in our study? We propose that *V. cholerae* uses the different blends of AIs it encounters along with environmental modulation of VqmA activity to gauge its changing locations in the host. Thus, VqmA functions rather like a “GPS-device”. In response to the information obtained about its micro-environment through VqmA, *V. cholerae* can appropriately tune its gene expression in space and time. We say this because, prior to entry into the small intestine (the site of cholera disease), *V. cholerae* will encounter oxygen limitation in the stomach. However, premature expression of virulence genes in the stomach, in the face of low pH and antimicrobial peptides would be unproductive and, moreover, divert energy from combating host defense systems. Thus, increased VqmA activity, due to enhanced accumulation of DPO under anoxia coupled with inhibition of formation of the activity-dampening intra-molecular disulfide bond, will increase production of VqmR and, in turn, repress expression of genes involved in biofilm formation and virulence. Anoxia and the concurrent presence of bile, encountered upon entry into the host duodenum (upper region of the small intestine), may provide a spatially relevant signal to alert *V. cholerae* to begin to express virulence genes. In this case, bile-mediated inhibition of VqmA activity due to prevention of formation of the inter-molecular disulfide bond will decrease VqmR production and, in turn, enhance expression of genes involved in biofilm formation and virulence. The combined use of bile and anoxia to decrease and increase VqmA activity, respectively, is also noteworthy because, as *V. cholerae* proceeds further through the intestinal tract, bile concentrations decrease near the ileum (lower portion of the small intestine), where ∼95% of bile salts are reabsorbed, and they reach a minimum in the large intestine. By contrast, anoxia is maintained throughout the intestinal tract. During successful infection, *V. cholerae* cell numbers increase as disease progresses. Accumulation of DPO should track with increasing cell density. Thus, it is possible that late in infection, *V. cholerae* resides in a high DPO, anoxic environment lacking bile, conditions enabling re-engagement of the VqmA-VqmR-DPO circuit, termination of virulence factor production, and expulsion from the host. In this context, we also note that DPO production by *V. cholerae* requires threonine as a substrate. Mucin is a major constituent of the intestinal tract and is composed of ∼35% threonine (26). Intriguingly, both the stomach and large intestine, locations where *V. cholerae* typically does not reside, contain more mucus secreting glands and mucus layers than does the small intestine (26). Thus, it is likely that enhanced access to mucus-derived threonine in the large intestine allows *V. cholerae* to increase DPO production. Again, high DPO levels repress virulence and biofilm formation. Thus, increased DPO production in the large intestine would foster *V. cholerae* departure from the host. We speculate that, in addition to oxygen and bile, the presence of mucus/threonine/the ability to synthesize DPO is also leveraged by *V. cholerae* as an additional spatial cue to optimize host dispersal timing.

## Materials and Methods

### Materials

iProof DNA polymerase was purchased from Biorad. Gel purification, plasmid-preparation, RNA-preparation (RNeasy), RNA-Protect reagents, qRT-PCR kits, and deoxynucleoside triphosphates were purchased from Qiagen. Antibodies were purchased from Sigma. Chitin flakes were supplied by Alfa Aesar. Instant Ocean (IO) Sea Salts came from from Tetra Fish. Bile salts were purchased from Fluka.

### Bacterial growth

*E. coli* Top10 was used for cloning, *E. coli* S17-1 λ*pir* was used for conjugations, and *E. coli* BL21 (DE3) was used for protein purification. *V. cholerae* and *E. coli* strains were grown in LB medium or in M9 minimal medium with glucose at 37°C, with shaking. When required, media were supplemented with streptomycin, 500 μg/mL; kanamycin 50 μg/mL; spectinomycin, 200 μg/mL, polymyxin B, 50 μg/mL; and chloramphenicol, 1 μg/mL. For bioluminescence assays, *V. cholerae* strains were cultured in SOC medium supplemented with tetracycline, and for AI-2 measurements, with boric acid (20 µM). Unless otherwise indicated, bile salts were supplemented at 0.5% v/v and DPO at 25 µM. Where indicated, oxygen deprivation was achieved as follows: a) medium was sparged prior to use for at least 20 min with nitrogen gas; b) medium was incubated overnight with constant stirring inside a COY anaerobic chamber equipped with a catalyst to scavenge oxygen. For both (a) and (b), subsequent steps were conducted inside the COY anaerobic chamber and c) exponentially growing cells were transferred into capped microcentrifuge tubes with a headspace to volume ratio (HV ratio) of zero and anaerobiosis was verified by the addition of 0.001% resazurin to control tubes, as described previously (18, 37). Micro-aerobiosis was achieved at ∼10 min post-shift and anaerobiosis by ∼15-18 min post-shift under these conditions, consistent with previous observations (18). Experiments presented in Figure 1 were conducted using strategy (a) and (b). Experiments in later sections used strategy (c). In control experiments, we verified that our data were not altered due to differences in growth in the presence or absence of oxygen (see Supplemental Figure 1B).

### Strain construction

#### Chromosomal alterations

*V. cholerae* strains were constructed using natural-transformation mediated multiplexed-genome-editing (MuGENT) (12, 13). Unless otherwise stated, chromosomal DNA from *V. cholerae* C6706 Sm^R^ was used as a template for PCR reactions. DNA fragments containing ∼3 kb of homology to the upstream and downstream regions of the desired chromosomal region were generated using PCR. When necessary, SOE PCR was used to combine multiple fragments of DNA, in which each fragment typically contained ∼27-30 bp of overhang homology (12, 13). Antibiotic resistance cassettes, to facilitate selection of transformants following the MuGENT step, were designed to integrate at a neutral locus (*vc1807*) and were gifts from the Dalia group (Indiana University) (12, 13). *V. cholerae* cultures for use in natural transformations were prepared by inoculating 1 mL liquid LB medium from freezer stocks and growing the cells to OD_600_∼1. Cells were pelleted at maximum speed in a micro-centrifuge and resuspended at the original volume in 1X IO Sea Salts (7 g/L). Competence was induced by combining a 75 μL aliquot of the cell suspension with 900 μL of a chitin IO mixture (8 g/L chitin), and the preparation was incubated overnight at 30°C. The next day, these mixtures were supplemented with one (or multiple) PCR-amplified linear DNA fragment(s) of interest, as well as DNA encoding an antibiotic resistance cassette (12, 13). These mixtures were incubated overnight at 30°C, followed by vortex for 10 min. Next, 150 μL of the suspension was plated onto solid LB medium containing the appropriate antibiotics followed by overnight incubation at 30°C. Resulting transformants were passaged three times on solid LB medium with antibiotics for purification. Genomic DNA from recombinant strains was used as a template for PCR to generate DNA fragments for future co-transformation, when necessary.

Site-directed mutations in *vqmA* were constructed by incorporating the desired alteration into forward or reverse PCR primers and generating DNA fragments with homology to DNA flanking chromosomal *vqmA*. These fragments were transformed, as described above, into a strain carrying *vmqA*::*kan* and the *vqmR-lacZ* transcriptional reporter integrated onto the chromosome. Clones were selected by screening for loss of kanamycin resistance and/or by assessment of positive LacZ activity when plated on agar containing 50 µg/mL X-gal.

#### Plasmid constructions

DNA cloned into the pBAD-pEVS or pEVS plasmids was assembled using enzyme-free XthA-dependent *in vivo* recombination cloning, as previously described (3, 41). Briefly, linear insert DNA fragments containing 30 base pairs of overlapping homology were generated using PCR. The plasmid backbone was likewise linearized by PCR-amplification. All DNA fragments were gel purified and eluted in ddH_2_O. Thereafter, 80 ng of the backbone and 240 ng of each insert DNA fragment were combined, incubated for 1 h at room temperature followed by transformation and clone recovery in chemically-competent Top10 *E. coli* cells. Constructs in the pET15b backbone were assembled using traditional restriction-enzyme cloning using primers and protocols described earlier (46).

### Assessing protein abundance and formation of disulfide bonds

Strains cultured overnight in LB medium (∼16-18 h) were diluted into fresh M9 minimal medium, with antibiotics, as necessary, to a final OD_600_=0.004. When assessing levels of VqmA-FLAG produced from the chromosomally-integrated *vqmA-FLAG* construct, strains were cultured to OD_600_∼0.3 (∼4 h of growth) and cells were harvested by centrifugation. For strains requiring induction of protein expression, 0.2% arabinose was added to the culture medium at 4 h post-inoculation and growth was continued for an additional 1 h at which point the cultures were divided and portions were supplemented with 25 µM DPO and/or 0.5% bile salts and/or deprived of oxygen. Treatments were continued for another 1 h after which cells were harvested by centrifugation at 13,000 rpm and the pellets were immediately frozen at -80°C until use.

### Immunoblotting

Cells were resuspended in ice-cold PBS and diluted to a final OD_600_=7 for protein produced from the chromosome or to OD_600=_3.5 for protein produced from a plasmid, in a volume of 20 µL. The cells were lysed by addition of 5 µL Bug Buster (Novagen) supplemented with 1 μg lysozyme, and 25 U/mL benzonase. Samples were combined with SDS-PAGE buffer in the presence or absence of DTT (100 mM), boiled for 20 min, and proteins separated on 4–20% Mini-Protein TGX gels (Biorad). Proteins were transferred to PVDF membranes (Bio-Rad) for 1 h at 4°C at 100 V. Membranes were blocked overnight in PBST (1X PBS, 0.03% Tween-20) supplemented with 5% milk, washed 5 times with PBST, and incubated for 40 min with 1:5000 dilution of monoclonal Anti-FLAG-Peroxidase antibody (Sigma) in PBST. The membranes were subsequently washed another five times with PBST. FLAG epitope-tagged protein levels were visualized using the Amersham ECL western blotting detection reagent (GE Healthcare). Thereafter, the antibody was removed by 2 serial incubations in stripping buffer (15 g/L glycine, 1 g/L SDS, 10 mL/L Tween-20, pH 2.2) for 5 min each. The membrane was re-equilibrated by 4 washes in PBST, 20 min each, and used to detect the abundance of the loading control, RNA Polymerase α. Washes and incubations as described above were performed to enable antibody binding and removal of excess. The primary antibody, anti-*E. coli* RNA Polymerase α (Biolegend), and the secondary antibody, anti-mouse IgG HRP conjugate antibody (Promega), were both used at a 1:10,000 dilutions. In all cases, protein levels were quantified using Image J software.

### Protein purification

pET15b plasmids encoding 6XHis-VqmA were mobilized into Δ*tdh E. coli* BL21 DE3. Strains were cultured for protein production as described previously (22, 46). Cells were harvested by centrifugation and pellets were resuspended in 1/100 volume of lysis buffer (50 mM Tris, 150 mM NaCl, pH 7.5 containing 0.5 mg/mL lysozyme, 1X protease inhibitor, and benzonase) for five min followed by the addition of an equal volume of Bugbuster Reagent (Novagen). The cell lysate was clarified by centrifugation at 13,000 rpm and protein was purified using Ni-NTA superflow resin (Qiagen), according to the manufacturer’s recommendations for a centrifugation-based protocol, except that the loading and wash buffers all contained 1- 5 mM imidazole to decrease non-specific protein binding. The protein was eluted from the resin using 300 mM imidazole and thereafter dialyzed twice against 50 mM Tris, 150 mM NaCl, pH 7.5, using a Slide-A-Lyzer module (Thermo Fisher). When necessary, buffers were amended with 5 mM DTT. To purify Holo-VqmA, buffers were supplemented with 100 µM DPO.

### Electromobility gel shift assays (EMSA)

The DNA corresponding to the promoter region of *vqmR*, ∼100 base pairs, was amplified using *V. cholerae* genomic DNA as a template. Where mentioned, protein was pre-treated in binding buffer with 10-fold molar excess of DTT or diamide in order to reduce or oxidize the protein, respectively. To initiate EMSA assays, 0.2-3.5 µM protein was combined with 30 ng probe DNA in binding buffer (50 mM Tris-HCl pH 8, 150 mM NaCl). Reactions were allowed to proceed at RT for 15 min. Samples were separated on a Novex 6% DNA Retardation Gel (Thermo) by electrophoresis in 1X TBE at 100 V. Gels were subsequently incubated with Sybr Green reagent, diluted in 1X TBE at RT for 25 min, washed with five successive rounds of ddH_2_O, and imaged using an ImageQuant LAS 4000 imager and the Sybr Green channel setting.

### Analysis of relative AI levels in conditioned growth medium following aerobic or anaerobic growth

*V. cholerae* strains were cultured overnight in LB medium (∼16-18 h), diluted into fresh aerobic or anaerobic LB medium to a final OD_600_=0.004. The cultures were incubated at 37°C for an additional ∼6 h with shaking. The cells were removed by centrifugation at 13,000 rpm and the spent medium filtered through 0.2 μm filters. 20% (v/v) 5X LB was added to the spent medium preparations (hereafter designated as reconditioned spent medium). Negative controls consisted of spent medium prepared from strains incapable of synthesizing CAI-1, AI-2, and DPO. Subsequently, reconditioned spent medium was combined with *V. cholerae* reporter strains expressing only a single QS receptor that, therefore detect only one AI. In the case of CAI-1 and AI-2 detection, the reporter strains carried a plasmid encoding the *V. harvyei luxCDABE* (luciferase) operon (39). For DPO detection, the reporter strain possessed a p*vqmR-lux* (luciferase) fusion on the chromosome in place of the native *lacZ* locus (22). Genotypes are provided in Table S1. The reporter strains and reconditioned spent medium preparations were combined to a final volume of 150 µL in wells of 96-well plates, covered with Breathe-Easy Film, and incubated at 30°C with shaking for 2.5-4 h. Finally, bioluminescence and OD_600_ values were recorded. Relative light units (RLU) were defined as light production (counts per minute) divided by OD_600_. Normalized RLUs were obtained by subtraction of the RLU values obtained from the negative controls.

### Beta-galactosidase assays

*V. cholerae* strains cultured overnight in LB medium were diluted into fresh M9 medium to a final OD_600_=0.004 and thereafter cultured to OD_600_∼0.3 (∼4 h). The cultures were held on ice prior to assay or subjected to further treatments, as described below. For strains requiring inducible protein expression, 0.01% arabinose was added to the culture medium at 4 h post-inoculation and growth was continued for an additional 1 h. In all cases, cultures were divided into aliquots and individual portions were supplemented with 25 µM DPO and/or 0.5% bile salts and/or deprived of oxygen. Cells were cultured for an additional 1 h and then held on ice prior to assay. To assess the effect of reductant, at 2 h post-inoculation, the cultures were supplemented with 300 µM DTT. Cells were cultured for another 2 h to allow DTT permeation, before *vqmA* expression was induced by the addition of arabinose. Subsequent treatments were as above. LacZ activity assays were carried out as follows: cells were combined 1:1 (v/v) with Bugbuster Reagent for 20 min. The assay was initiated by combining 20 µL of the cell/Bugbuster mixture with 140 µL of assay buffer (80% Bugbuster, 10% 10X PBS, 1 mM MgSO_4_, 10 µg/mL lysozyme, benzonase (0.05% v/v), β-mercaptoethanol (0.1% v/v), 67 µg/mL fluorescein-di-β-D-galactopyranoside). Changes in fluorescence were captured using the GFP channel on a Synergy Neo2 HTS Multi-Mode Microplate Reader. Activity units were defined as the change in fluorescence/minute/ OD_600_ of the culture at the point of harvest.

### RNA isolation and quantitative RT-PCR

Strains cultured overnight in LB were diluted into fresh M9 minimal medium to a final OD_600_=0.004. Next, the strains were grown to OD_600_∼0.3 (∼4 h post-inoculation) with shaking at 37°C at which point the cultures were divided into portions that were supplemented with 25 µM DPO and/or 0.5% bile salts and/or deprived of oxygen. Treatments were continued for another 1 h, the cells were harvested and treated for 15 min at room temperature with RNAProtect reagent, as per the manufacturer’s instructions. RNA was isolated using the RNeasy kit (Qiagen) and 2 µg of total RNA was depleted of contaminating DNA using TurboDNase (Applied Biosystems), using the manufacturer’s recommended protocol. 500 ng of the resulting total RNA was used to construct cDNA libraries using SuperScript III Reverse Transcriptase (Invitrogen). q-PCR was conducted using the PerfeCTa SYBR Green FastMix Low ROX (Quanta Biociences) reagent.

### Caco-2 culture and co-culture with bile and *V. cholera*

The HTB37 cell line was obtained from ATCC and thereafter cultured and passaged in EMEM (ATCC) supplemented with an 10% FBS (Thermo Fisher), 2 mM glutamine (Thermo Fisher), 1X Penstrep (Thermo Fisher) and 2.5 µg/mL plasmomycin (Invitrogen), as per ATTC recommendations. Prior to co-culture, Caco-2 cells were seeded at 0.32 cm^2^ into a tissue culture treated 96-well plate in EMEM medium, as above, except that the 2 mM glutamine was not included. The cells were cultured to confluency, the medium was removed by aspiration, and the cells were washed with Earle’s balanced salt solution followed by the addition of EMEM medium containing 25 µM DPO, but lacking antibiotics/glutamine/FBS. At time zero, *V. cholerae*, grown as described below, was added, at an MOI of 10 and with 0.05% bile salts. Following co-culture for 3.5 h, the medium and the bacteria were removed by aspiration, the wells were washed with Earle’s balanced salt solution, and Caco-2 cell viability was assessed using the neutral red assay (51). For co-culture with Caco-2 cells, *V. cholerae* WT and Δ*vqmA* strains were cultured overnight under static conditions in AKI medium containing 25 µM DPO in tubes with an HV ratio of zero at 37°C. The next day, the cells were decanted into glass tubes and cultured with vigrous shaking for 1 h. Subsequently, the cultures were diluted with PBS to the appropriate density and added to the Caco-2 cells.

## Acknowledgements

We thank the Donia lab for generously allowing us to use their anaerobic chamber. We thank Dr. Ankur Dalia for the gift of protocols, strains, and reagents to facilitate MuGENT cloning. We thank Dr. Tharan Srikumar, Saw Kyin, and Henry Shwe for help with mass spectroscopy experiments. This work was supported by the Howard Hughes Medical Institute, National Science foundation Grant MCB- 1713731, and NIH Grant 5R37GM065859 to B.L.B. A.A.M is a Howard Hughes Medical Institute Fellow of the Life Sciences Research Institute.

