## Supplemental Data for "The *Vibrio cholerae* quorum-sensing protein VqmA integrates cell density, environmental, and host-derived cues into the control of virulence"

### Supplementary Information

#### Supplementary Methods:

**Analysis of CAI-1 levels following shifts in oxygen:** *V. cholerae* strains grown overnight at 37°C were diluted into fresh LB medium and incubated at 37°C with shaking to $OD_{600}$>1 under aerobic conditions. Next, the cells were removed by centrifugation for 1 min at 13,000 rpm and a portion of the spent medium was collected. This preparation is referred to as 'Pre' in Figure S1B. The cell pellets were resuspended in equal volumes of fresh aerobic or anaerobic LB medium. A portion of the added medium was immediately removed and saved. This preparation serves as the time 0 sample in Figure S1B. The cultures were subsequently incubated in the presence or absence of oxygen, and spent media preparations were generated periodically as above. Following collection, the spent media were filtered through 0.2 µm filters. 20% (v/v) 5X LB was added to the preparations (hereafter designated as reconditioned spent medium). Subsequently, reconditioned spent medium was combined with a *V. cholerae* reporter strain that detects only exogenously supplied CAI-1 and carries a plasmid encoding the *V. harvyei luxCDABE* genes (1).

**Mass Spectrometry Data Acquisition.** Purified 6XHis-VqmA was treated with diamide followed by desalting using a C8 zip tip, dried using a SpeedVac, and dissolved in 155 µL of 50 mM ammonium bicarbonate buffer. Thereafter, accessible cysteine residues were alkylated by incubation in the presence of 18 mM CAA, at room temperature. One half of the sample was retained and subjected to mass spectrometry analysis (below). The second half of the sample was desalted, dried, and resuspended in ammonium bicarbonate buffer and cysteine residues were reduced by treatment with 10 mM TCEP and desalted. Subsequently, alkylation was achieved by incubation in the presence of 18 mM NEM. Digestion was accomplished by incubation with 500 ng of endoproteinase Glu-C overnight at 37°C.

Digested samples were dried completely in a SpeedVac and resuspended with 21 µL of 0.1% formic acid, pH 3. Thereafter, 2 µL of the samples were injected, using an Easy-nLC 1200 UPLC system, onto a 1.9 µm C18-AQ nano capillary column (Dr. Maisch, Germany; 45 cm long with 75 µm inner diameter). The columns were mated to a metal emitter in-line with an Orbitrap Fusion Lumos (Thermo Scientific, USA). Samples were resolved using a 1 h gradient (300nL/min flow rate; $45^{o}$C column temperature). The mass spectrometer was operated in data dependent mode with the MS1 scan conducted at a resolution of 120,000 and the following settings: positive mode, profile data type, AGC $4 \times 10^5$, Max IT 54 ms, 300-1500 m/z. This procedure was followed by high energy collision dissociation (HCD) fragmentation with 35% collision energy. A dynamic exclusion list was invoked for 60 s with a maximum cycle time of 3 s to exclude previously fragmented peptides. Peptides were isolated for fragmentation using a quadrupole with a 1.2 Da window.

**Mass spectrometry data analysis.** Raw files were analyzed using the PEAKS Studio software (v. 10.0; Bioinformatics Solutions Inc) (2, 3). Parent ion and fragment tolerance were set at 70 ppm and and 0.100 Da, respectively. The oxidation of methionine residues, acetylation of protein N-termini, N-ethylmaleimide modification of cysteine residues, the conversion of glutamine residues to pyro-glutamate, and the deamidation of asparagine residues were specified as dynamic modifications. Files were searched against a database containing the His6-VqmA sequence. The VqmA sequence was obtained from the GPM database: ftp://ftp.thegpm.org/fasta/cRAP.

The Scaffold software (v. 4.8.4, Proteome Software Inc., Portland, OR) was used to validate MS/MS-based peptide and protein identifications. Peptide identification was conducted using the Scaffold Local FDR algorithm (cutoff of >90.0%). Protein probabilities were assigned using the Protein Prophet algorithm with a cutoff of >99% and a minimal requirement of least two matching peptide reads.

**Supplementary Figures and Figure Legends:**

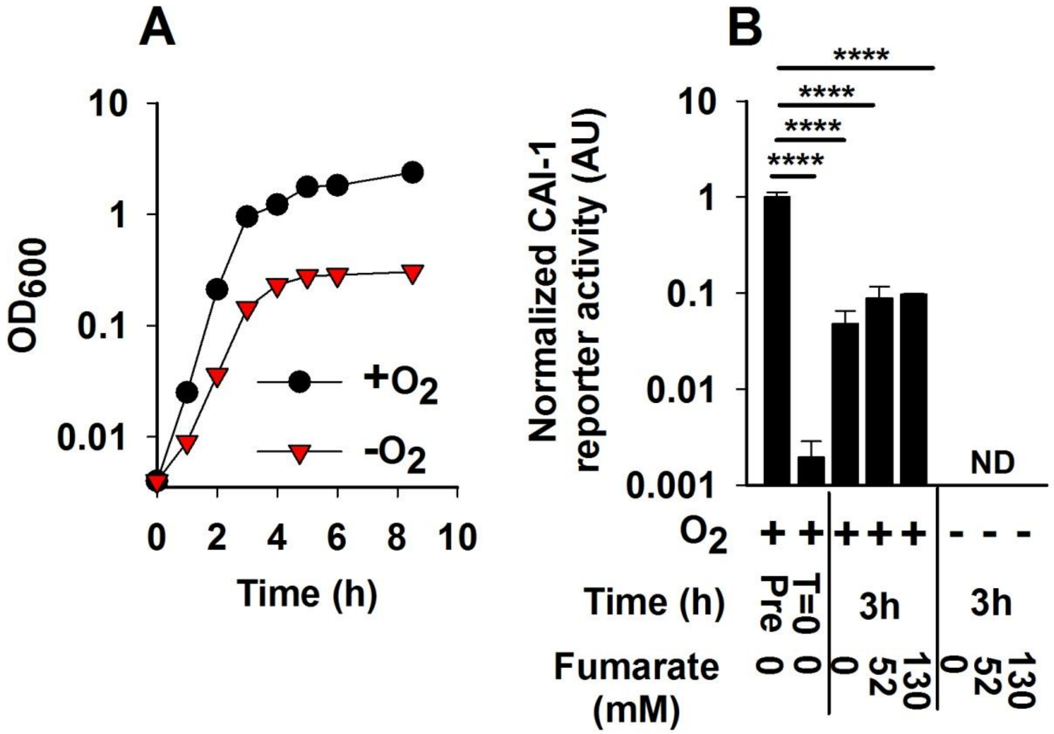


**Figure S1.** The absence of oxygen reduces *V. cholerae* growth and CAI-1 is not produced under anaerobic conditions. **(A)** Growth of WT *V. cholerae* in the presence (black) and absence (red) of $O_2$ in LB medium. **(B)** 80% cell-free culture fluids were were provided to a *V. cholerae* reporter strain that produces bioluminescence in response to exogenous CAI-1. The fluids were prepared from WT *V. cholerae* cultured aerobically to high cell density (Pre), a point when CAI-1 production is maximal, followed by washing and resuspension +/- $O_2$ and/or +/- fumarate for 3 h. This strategy ensured that an equal number of CAI-1-producing cells with an equivalent capacity to synthesize CAI-1 were present when oxygen was removed and/or fumarate was provided. CAI-1 activity immediately following resuspension is designated T=0. Data in panels A and B represent the average values of biological replicates (*n*=3) and error bars represent SD. **** denotes $p<0.0001$. ND denotes that levels of CAI-1 were below the level of detection in our assay.

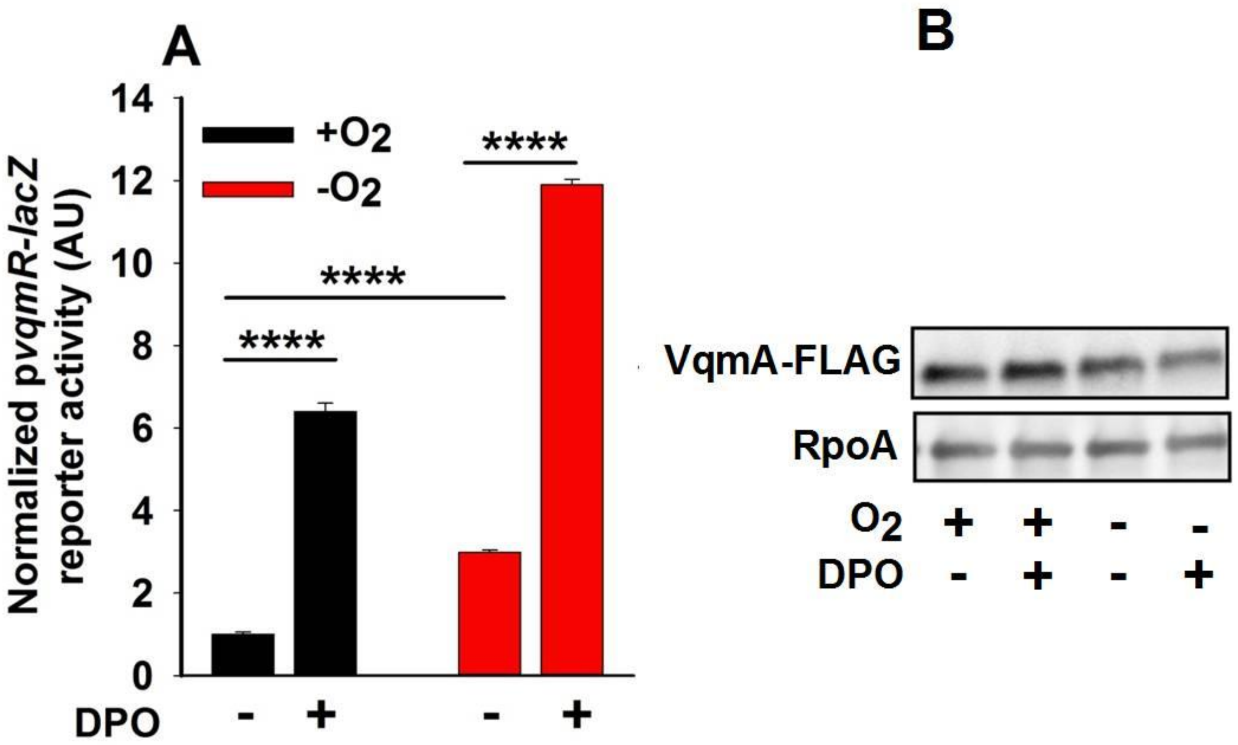


**Figure S2.** The absence of oxygen enhances the activity of endogenously expressed Apo- and Holo-VqmA-FLAG and these phenotypes are not due to altered levels of VqmA-FLAG. **(A)** Relative p*vqmR-lacZ* expression and **(B)** Western blot showing VqmA-FLAG abundance with RpoA abundance as the loading control in a Δ*tdh V. cholerae* strain carrying *vqmA-flag* at its endogenous location in the chromosome. The cells were exposed for 1 h to -$O_2$, +$O_2$, and/or 25 µM DPO. Data in (A) represent the average values of biological replicates (*n*=3) and error bars represent SD. **** denotes $p<0.0001$.

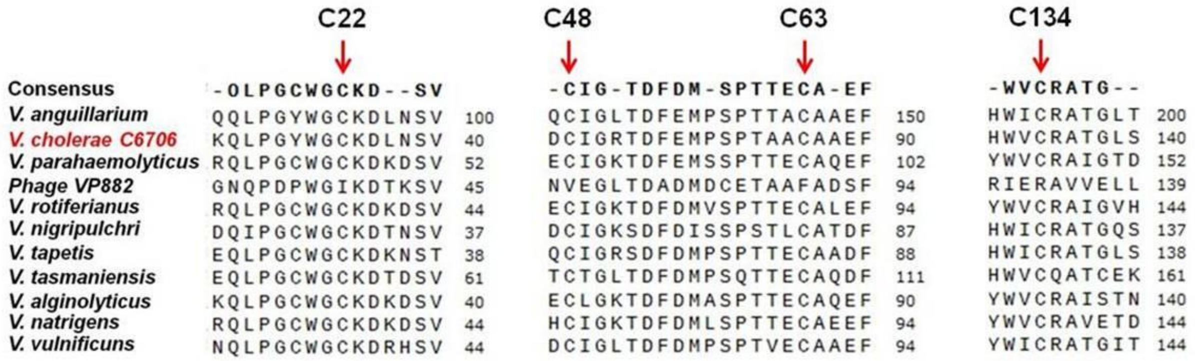


**Figure S3.** Cysteine residues in VqmA are conserved across homologs from *Vibrionaceae* but not in a VqmA-containing vibriophage. Multiple sequence (SnapGene) alignments are displayed for select regions in the VqmA polypeptide sequences.

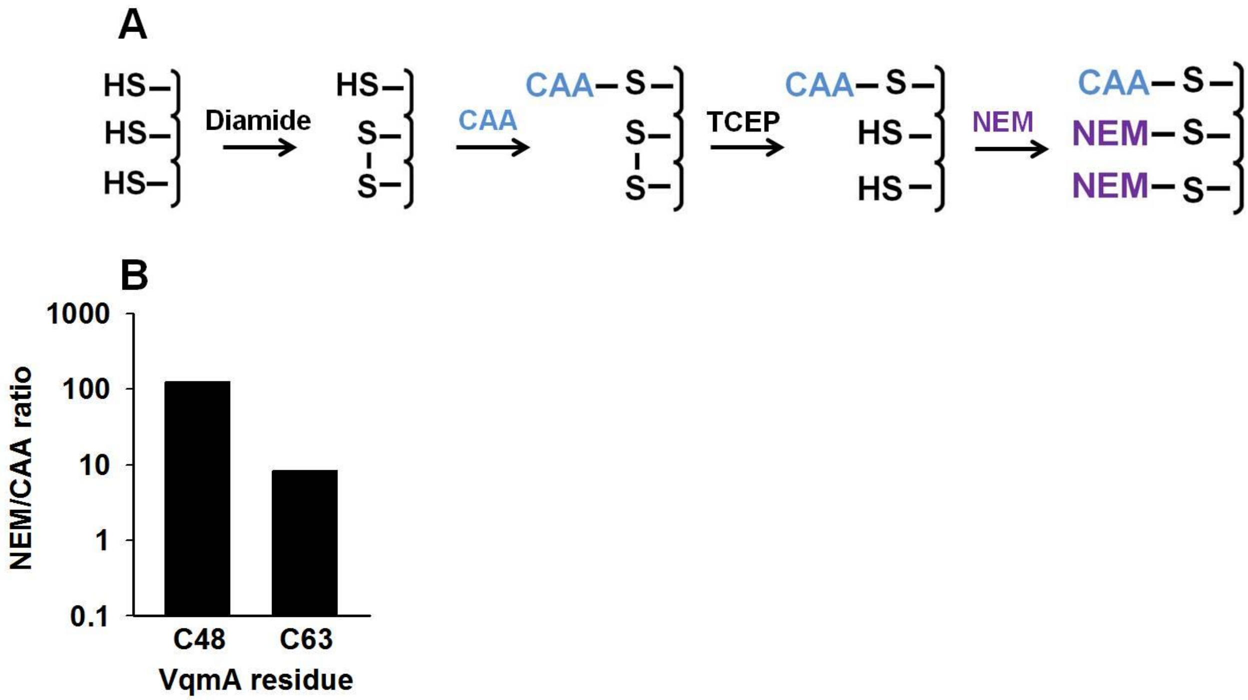


**Figure S4.** Thiol trapping analyses suggest that VqmA C48 and C63 are involved in disulfide bonds. **(A)** Schematic for the thiol trapping strategy with chloroacetamide (CAA) and N-ethylmaleimide (NEM). **(B)** Quantitation of mass spectral counts for C48 and C63 following thiol trapping with NEM and CAA. 6His-VqmA, oxidized by incubation with diamide was denatured and incubated with CAA. The sample was desalted, the protein was reduced with TCEP, and again desalted. The protein was next incubated with NEM. Samples were proteolyzed and subjected to mass-spectrometry. The data displayed represent the average of three separate injections.

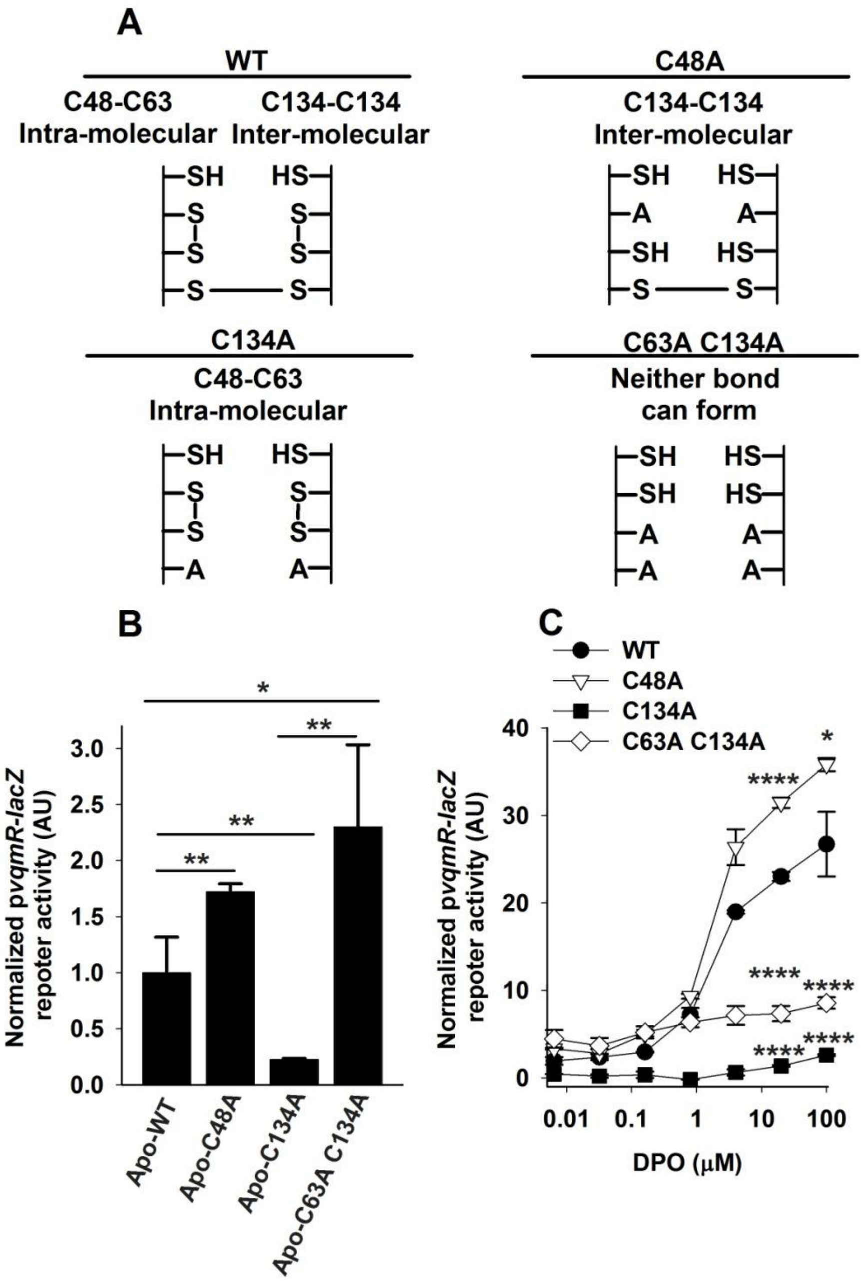


**Figure S5.** VqmA activity is modulated by intra- and inter-molecular disulfide bonds. **(A)** Schematic displaying disulfide bond formation in the VqmA variants studied here. **(B and C)** p*vqmR-lacZ* activity in the $\Delta$*tdh V. cholerae* strain carrying the designated *vqmA-FLAG* alleles expressed from the native chromosomal location following growth in the presence of $O_2$ and without **(B)** or with **(C)** DPO. Data represent the average values of biological replicates (*n*=3) and error bars represent SD. * denotes p<0.05, ** denotes p<0.01, and **** denotes p<0.0001.

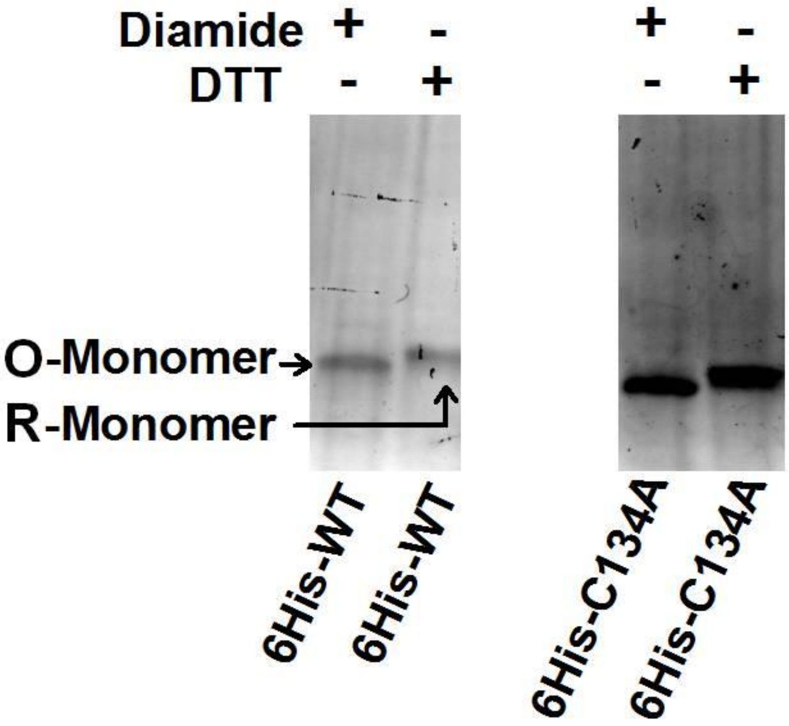


**Figure S6.** Purified VqmA protein forms the C48-C63 intra-molecular disulfide bond. SDS-PAGE of the designated purified 6His-VqmA proteins. Treatment with 10-fold molar excess diamide or DTT as indicated.

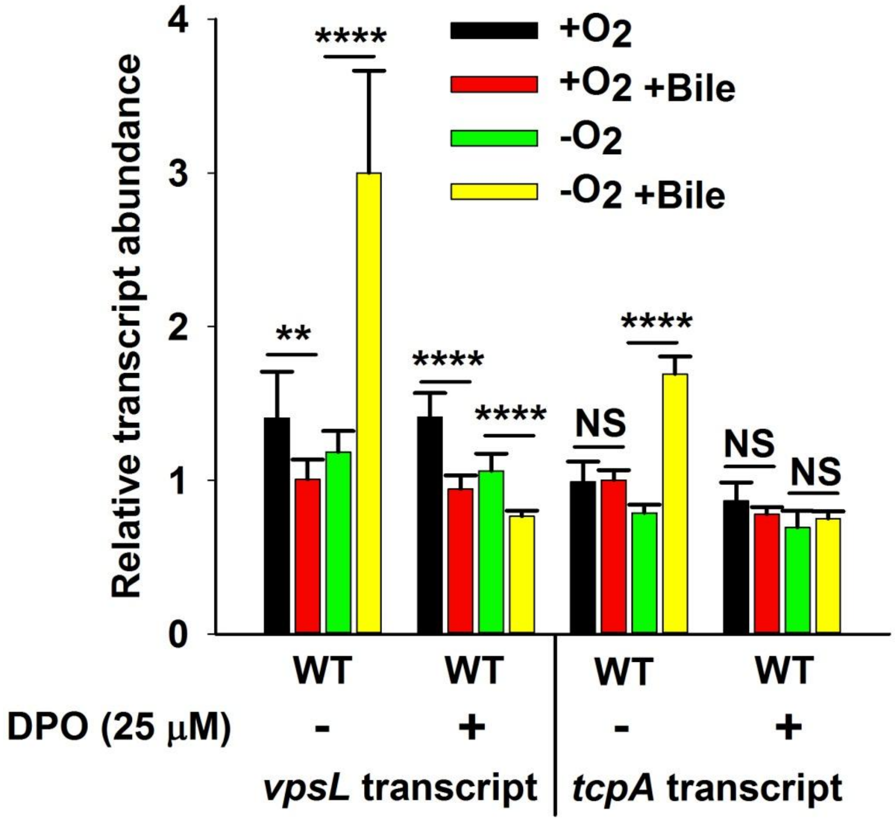


**Figure S7.** Bile drives increased *vpsL* and *tcpA* expression exclusively in the absence of oxygen. Relative expression levels of *vpsL* and *tcpA* in WT *V. cholerae* treated as specified. ** denotes $p<0.01$, **** denotes $p<0.0001$, and NS denotes $p>0.05$.

### Supplementary Tables:

**Table S1. Strains and plasmids used in this study.**

**Strains used in this study**

| **Strains** | **Genotype/Description** | **Genetic Background** | **Source/Reference** |
|---|---|---|---|
| BB-VC 90 | *Vibrio cholerae*; Wild-type; C6706; Streptomycin resistant | C6706 | Bassler Lab Collection |
| BB-VC 12 | Δ*vqmA* | C6706 | Bassler Lab Collection |
| BB-VC 38 | Δ*tdh* | C6706 | Bassler Lab Collection |
| BB-VC 8 | Δ*cqsA* | C6706 | Bassler Lab Collection |
| BB-VC 90 | Δ*luxS* | C6706 | Bassler Lab Collection |
| BB-VC 176 | Δ*cqsS* Δ*vpsS* Δ*cqsR* Δ*luxS* | C6706 | Bassler Lab Collection |
| BB-VC 0325 | Δ*tdh lacZ::pvqmR-lux* | C6706 | Bassler Lab Collection |
| AM 36 | Δ*cqsR* Δ*vpsS* Δ*luxQ* Δ*cqsA* | C6706 | Bassler Lab Collection |
| AM3 | Δ*tdh lacZ::pvqmR-lacZ* | C6706 | This work |
| AM565 | Δ*tdh vqmA::kan lacZ::pvqmR-lacZ VC1807::spec* | C6706 | This work |
| AM616 | Δ*tdh* Δ*vqmA lacZ::pvqmR-lacZ VC1807::Cm* | C6706 | This work |
| AM629 | Δ*tdh vqmA-flag lacZ::pvqmR-lacZ VC1807::spec* | C6706 | This work |
| AM625 | Δ*tdh vqmA-C22A flag lacZ::pvqmR-lacZ VC1807::spec* | C6706 | This work |
| AM626 | Δ*tdh vqmA-C48A flag lacZ::pvqmR-lacZ VC1807::spec* | C6706 | This work |
| AM627 | Δ*tdh vqmA-C63A flag lacZ::pvqmR-lacZ VC1807::spec* | C6706 | This work |
| AM628 | Δ*tdh vqmA-C134A flag lacZ::pvqmR-lacZ VC1807::spec* | C6706 | This work |
| AM421 | *Escherichia coli* | BL21 | Bassler Lab Collection |
| BB-EC | Δ*tdh* | BL21 | Bassler Lab Collection |

**Plasmids used in this study**

| **Plasmid name** | **Insert Locus/function** | **Source/Reference** |
|---|---|---|
| pEVS-pBAD | Cloning vector for arabinose inducible expression | Bassler Lab Collection |
| pEVS-pBAD-*vqmA flag* | Inducible expression & western blots | This work |
| pEVS-pBAD-*vqmA*-C22A *flag* | Inducible expression & western blots | This work |
| pEVS-pBAD-*vqmA*-C48A *flag* | Inducible expression & western blots | This work |
| pEVS-pBAD-*vqmA*-C63A *flag* | Inducible expression & western blots | This work |
| pEVS-pBAD-*vqmA*-C134A *flag* | Inducible expression & western blots | This work |
| pEVS-pBAD-*vqmA*-C63A C134A *flag* | Inducible expression & western blots | This work |
| pET28b | Cloning vector for protein production | Bassler Lab Collection |
| pET28b-*his6-vqmA* | Protein production in *E. coli* | This work |
| pET28b-*his6-vqmA C134A* | Protein production in *E. coli* | This work |
| pBB1 | Luciferase-based quorum sensing reporter | Bassler Lab Collection |

**Table S2. Oligonucleotides used in this study.**

| Number | Oligonucleotide | Sequence |
|---|---|---|
| **Cloning** | | |
| AM7 | VCA1078_Mugent_For | ATTGAAATGCGTCTGTCGCAAATCAAACAGCG |
| AM8 | VCA1078_Mugent_Rev | AAACACCGCAATCATCCCTGCGACTAG |
| AM9 | VC1807_Mugent_For | TTTAAAGGGGATCAGTGACCG |
| AM10 | VC1807_Mugent_For | CAATTTTGCTTTTGGACCATCCC |
| AM41 | VCA1078_C22A_For | ATTACCCGGTTATTGGGGAGCCAAGGACTTAAACTCGG |
| AM42 | VCA1078_C22A_Rev | CCGAGTTTAAGTCCTTGGCTCCCCAATAACCGGGTAAT |
| AM43 | VCA1078_C48_Rev | AAAATCGGTGCGCCCGATGGCATCTTCAGCGCGCTTTAAG |
| AM44 | VCA1078_C48_For | CTTAAAGCGCGCTGAAGATGCCATCGGGCGCACCGATTTT |
| AM45 | VCA1078_C63_For | GCCTAGCCCAACAGCAGCTGCCGCTGCCGAATTTCAACAG |
| AM46 | VCA1078_C63_Rev | CTGTTGAAATTCGGCAGCGGCAGCTGCTGTTGGGCTAGGC |
| AM47 | VCA1078_C134_For | TGAAGTTGGTCATTGGGTCGCCCGAGCAACTGGGTTATCC |
| AM48 | VCA1078_C134_Rev | GGATAACCCAGTTGCTCGGGCGACCCAATGACCAACTTCA |
| AM269 | pBAD_up_ For | ACTGTTTCTCCGGATCCAAGGAGTGTATTCGTGCCTAACCATCTGA CATTAGAGCAGAT |
| AM270 | pBAD_up-rev | ATCTGCTCTAATGTCAGATGGTTAGGCACGAATACACTCCTTGGAT CCGGAGAAACAGT |
| AM271 | pBAD_down_for | TTGATTGGGCTTATGGCGCCAGTCTATCGAGGATCCGGTGATTGA TTGAGCAAGCTTTA |
| AM272 | pBAD_down_rev | TAAAGCTTGCTCAATCAATCACCGGATCCTCGATAGACTGGCGCC ATAAGCCCAATCAA |
| **Electromobility Gel Shift Analyses** | | |
| AM353 | | TGTTGACTCAAACAATTATGCA |
| AM360 | | GGTTTGTACTTTACCGAACGCGGTA |
| **Quantitative Real-Time PCR Analyses** | | |
| AM-RT-1 | VC1258_*gyr*_RT_For | TGGCCAGCCAGAGATCAAG |
| AM-RT-1 | VC1258_ *gyr*_RT_Rev | ACCCGCAGCGGTACGAT |
| AM-RT-5 | VC0934_ *vpsL*_RT_For | CAGTATGCGAGTGATGGATAATGG |
| AM-RT-6 | VC0934_*vpsL*_RT_Rev | TCGTGGATCGCCTTTGGT |
| AM-RT-57 | VC0828_ *tcpA*_RT_For | GTGGTCTCAGCGGGTGTTGTTAC |
| AM-RT-58 | VC0828_*tcpA*_RT_Rev | CCAAGACTACGATAAGTTTGTGTCATTGC |
